# Extensive transcriptional and chromatin changes underlie astrocyte maturation *in vivo* and in culture

**DOI:** 10.1101/2020.04.28.066043

**Authors:** Michael Lattke, Robert Goldstone, Francois Guillemot

**Author notes:** **Author List Footnotes** lead author (Francois Guillemot).

## Abstract

Astrocytes have diverse functions in brain homeostasis. Many of these functions are acquired during late stages of differentiation when astrocytes become fully mature. The mechanisms underlying astrocyte maturation are not well understood. Here we identified extensive transcriptional changes that occur during astrocyte maturation and are accompanied by chromatin remodelling at enhancer elements. Investigating astrocyte maturation in a cell culture model revealed that *in vitro*-differentiated astrocytes lacked expression of many mature astrocyte-specific genes, including genes for the transcription factors Rorb, Dbx2, Lhx2 and Fezf2. Forced expression of these factors *in vitro* induced distinct sets of mature astrocytes-specific transcripts. Culturing astrocytes with FGF2 in a three-dimensional gel induced expression of *Rorb*, *Dbx2* and *Lhx2* and improved their maturity based on transcriptional and chromatin profiles. Therefore extrinsic signals orchestrate the expression of multiple intrinsic regulators, which in turn induce in a modular manner the transcriptional and chromatin changes underlying astrocyte maturation.

## Introduction

Astrocytes are the most abundant glial cells in the mammalian central nervous system and they serve essential functions in brain development and homeostasis. Astrocytes are generated from radial glia, the principal neural stem cells (NSCs) of the brain, during late embryonic and early postnatal stages. First, a “gliogenic switch” enables NSCs to progress from neurogenesis to the generation of astrocytes and oligodendrocytes. Specified glial precursors then migrate from the progenitor zones to the brain parenchyma where they proliferate and differentiate during early postnatal stages (Ge et al., 2012; Molofsky and Deneen, 2015). During a subsequent phase of maturation in the first few postnatal weeks, immature astrocytes exit the cell cycle and aquire a fully mature phenotype (Molofsky and Deneen, 2015).

The molecular mechanisms controlling astrocyte specification and the early steps of differentiation of immature astrocytes have been intensely investigated and diverse regulators of these processes have been identified, including transcription factors such as NFIA and Sox9 and extrinsic signals such as bone morphogenetic proteins (BMPs) and JAK-STAT pathway ligands (Deneen et al., 2006; He et al., 2005; Kang et al., 2012; Kohyama et al., 2010). In contrast, little is known of the mechanisms controlling the later step of maturation of postnatal immature astrocytes into fully mature adult astrocytes.

Postnatal maturation is associated with major changes in astrocyte biology. Before their maturation, astrocytes contribute to brain vascularisation, establishment of the blood-brain-barrier, axon pathfinding and establishment and elimination of synapses (Allen and Lyons, 2018). These functions are specific of immature astrocytes and the genes involved, such as *Tnc* and *Sparc*, are downregulated when astrocytes mature (Dallerac et al., 2018). Conversely, maturing astrocytes acquire new functions required for adult brain homeostasis. They provide metabolic and trophic support for neurons, regulate the blood-brain-barrier and local cerebral blood flow, and modulate neuronal activity. Genes involved in these functions, including the glutamate transporter *Scl1a2*/*GLT-1*, connexins such as *Gjb6*/*Cx30*, and *Aqp4* are induced during postnatal maturation (Allen and Lyons, 2018; Dallerac et al., 2018).

Although the functions of astrocytes change profoundly during their maturation, the regulatory mechanisms underlying these changes have only recently been examined. This analysis has been performed using mostly in vitro models, including cultures of acutely isolated postnatal astrocytes, cultures of NSCs differentiated with agonists of the BMP or JAK-STAT signalling pathways (Bonaguidi et al., 2005; Kleiderman et al., 2015; Krencik et al., 2011; Tiwari et al., 2018), and pluripotent stem cell-derived three dimensional neural cultures or spheroids, (Sloan et al., 2017). Several mechanisms have been shown to promote maturation-associated transcriptional changes in these cell culture models, including the factors Runx2 and p53, the FGF signaling pathway and cell-cell-contacts (Li et al., 2019; Roybon et al., 2013; Tiwari et al., 2018). However, *in vitro*-differentiated astrocytes are thought to remain immature (Hasel et al., 2017; Roybon et al., 2013; Yang et al., 2013), and it is therefore unclear whether the mechanisms identified are sufficient to account for the complete maturation of astrocytes *in vivo*.

To gain insights into the mechanisms driving astrocyte maturation, we have characterised acutely isolated murine astrocytes at both postnatal and adult stages. Single-cell RNA sequencing revealed a progression of maturing astrocytes through distinct transitional stages. By integrating chromatin accessibility and gene expression data, we found extensive chromatin remodelling associated with the transcriptional changes in maturing astrocytes, which led us to identify potential regulators of *in vivo* maturation. We also studied astrocytes differentiated in vitro and confirmed that these cells do not become fully mature. Reconstitution experiments showed that transduction of the transcription factors Rorb, Fezf2, Dbx2 and Lhx2 in *in vitro*-differentiated astrocytes induced distinct maturation-associated gene modules. Moreover addition of FGF2 and cell-cell-contact-dependent signals to these cultures promoted the expression of maturation-inducing transcription factors as well as global transcriptional and chromatin changes associated with astrocyte maturation *in vivo*. Overall our results suggest that astrocyte maturation is promoted by extrinsic signals that induce multiple transcription factors that act largely independently to regulate distinct gene expression modules that together promote a mature astrocytic phenotype.

## Results

### Astrocyte maturation *in vivo* proceeds through transitional stages

To gain a better understanding of the molecular mechanisms underlying astrocyte maturation, we examined astrocytes isolated from mice at early postnatal stages (postnatal days 3 and 7; P3/P7) when astrocytes have been specified but remain immature and partially proliferating, and at young adult stages (~3 months) when astrocytes have reached full maturity (Fig. 1A). We collected tissues from the early postnatal and adult striatum, which we dissociated and enriched for astrocytes using magnetic-activated cell sorting (MACS) with the astrocyte surface antigen ACSA2 (Kantzer et al., 2017), and performed single cell RNA-Sequencing (scRNA-Seq) (Fig. 1B). We sequenced a total of 2671 cells from two independent preparations each of early postnatal and adult tissues. To identify cell types, we performed dimension reduction with the Uniform Manifold Approximation and Projection (UMAP) approach implemented in Seurat 3. This analysis revealed 23 cell clusters that we defined using a panel of established lineage markers (Figure 1C, S1A/B). 1391 astroglial lineage cells, identified by expression of Sox9 (Sun et al., 2017) and other well-established early and late astrocyte markers, separated into several clusters (Figure 1C, S1A/B). We further examined the expression of proliferation markers and the age of the mice from which the cells originated (Figure 1D), to determine which clusters represent immature or mature stages of astrocytic development.

**Figure 1:**
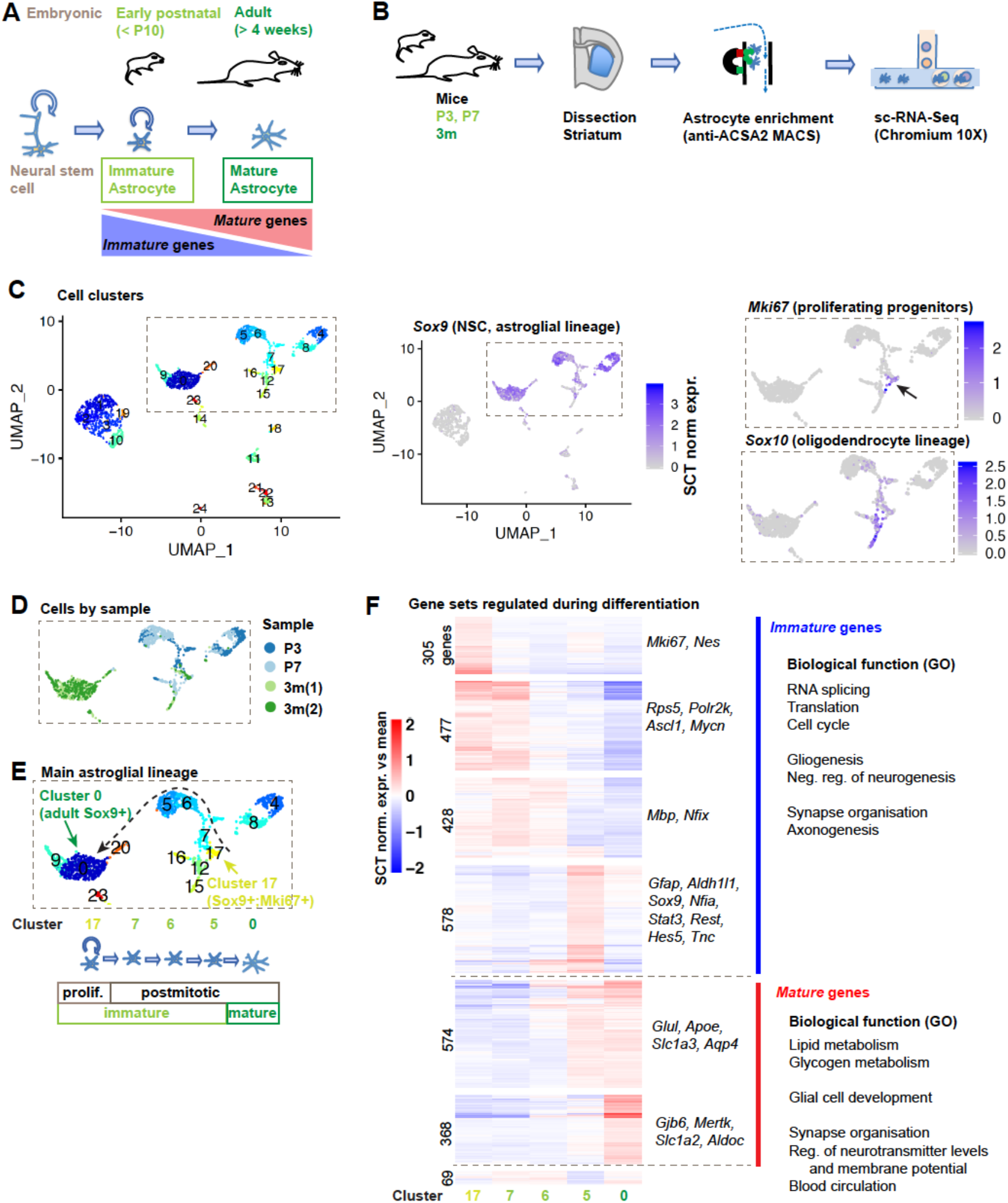
Astrocyte maturation *in vivo* proceeds through intermediate stages. **(A)** Current model of astrocyte differentiation in mice: neural stem cells generate immature astrocytes that proliferate until around postnatal day 10 (P10) and become fully mature astrocytes by 4 weeks of age. **(B)** Approach for single cell RNA-Seq analysis of striatal astrocyte maturation: immature astrocytes from the striatum of P3/P7 mice and mature astrocytes from the striatum of 3-months-old mice are enriched by magnetic bead assisted cell sorting (MACS) for the astrocyte-specific ACSA2 antigen, followed by scRNA-Seq. **(C)** UMAP dimension reduction plots of combined single cell transcriptomes from postnatal and adult striatum. Expression of *Sox9*, *Mki67* and *Sox10* is projected on UMAP plots. See also Figure S1. **(D)** Origin of *Sox9*+ cells highlighted in (C) by sample and developmental stage. **(E)** Hypothetical astrocyte differentiation trajectory deduced from (C) and (D): *Sox9*+ *Mki67*+ putative proliferating glial progenitors (cluster 17) differentiate into the main adult astrocyte cluster 0 via the early postnatal *Sox9*+ *Mki67*-clusters 7/6/5 (postmitotic immature astrocytes). **(F)** Differentially expressed genes along the hypothetical differentiation trajectory shown in (E); heatmaps show the mean relative expression of each gene in each cell cluster; genes were ordered by hierarchical clustering and grouped into gene sets with similar expression patterns. Example genes and potential biological functions based on gene ontology (GO) analysis are shown. 2799 differentially expressed genes were detected with Seurat3 by comparing the 900 cells of clusters 0, 5, 6, 7, 17. See also Figure S1, Table S1.

The astroglial clusters included several postmitotic populations (i.e. that did not express cell cycle genes) present only in the postnatal preparation (clusters 4-8) and one major cluster present only in the adult preparation (cluster 0), as well as three additional small populations of adult-only astrocytes (clusters 9, 20, 23; Fig. 1C,D). A small population (cluster 17) expressed proliferation markers including Mki67, as well as low levels of both astroglial and oligodendroglial markers including Sox9 and Sox10, respectively (Figure 1C, S1A,B). We hypothesise that these cells represent proliferating bipotent glial progenitors at the beginning of the astrocyte differentiation trajectory. This trajectory likely ends with the main adult astrocyte cluster 0 representing the fully mature state. The locations of the early postnatal clusters 7, 6 and 5 on the UMAP plot suggest that they represent consecutive intermediate states of astrocytic differentiation between clusters 17 and 0 (Figure 1C-E).

To gain insights into the process of astrocyte maturation, we analysed gene expression differences along this hypothetical differentiation trajectory (Figure 1E). Using Seurat, we identified 2799 genes that were differentially regulated between any of the clusters. To characterise their expression patterns across the different cell populations, we performed hierarchical clustering based on average expression of each gene in each cell cluster. We grouped the genes with similar expression patterns into six main gene sets (Figure 1F, Table S1). Four of these gene sets contained 1788 immature striatal astrocyte genes expressed at highest levels in one or several early postnatal astrocyte clusters (cluster 17, 7, 6 or 5), The remaining two gene sets contain 942 mature striatal astrocyte genes, most highly expressed in the mature adult astrocyte cluster 0 (Figure 1F, Table S1).

We next investigated the potential biological functions of these immature and mature genes, using the literature and Gene Ontology analysis (Figure 1F, Table S1). The immature gene sets included many genes of the basal transcription and translation machineries and genes involved in cell proliferation, e.g. the regulators of NSC proliferation *Ascl1* and *Mycn*. Immature genes were also enriched for GO terms linked to neuronal and glial development such as “negative regulation of neurogenesis“, “gliogenesis” and “axonogenesis” and included transcription factors involved in astrocyte differentiation such as *Sox9*, *Nfia*, *Stat3*, *Hes5* and *Rest*. In contrast, the mature genes were enriched for GO terms linked to lipid metabolism and terms related to homeostatic support functions of astrocytes, such as “synapse organization”, “regulation of neurotransmitter levels” and “blood circulation” and included many genes known to be involved in these functions, such as *Glul* (glutamine synthetase), *Gjb6* (*Cx30*), *Aldoc, Apoe*, *Aqp4* and the glial glutamate transporters *Slc1a3* (*Glast*) and *Slc1a2* (*GLT-1*) (Figure 1F, Table S1).

Overall, these transcriptional changes support the hypothetical astrocyte differentiation trajectory shown in Figure 1E and suggest that proliferating and bipotent progenitors differentiate to fully mature astrocytes through at least three intermediate stages that are postmitotic but still immature. The gene expression analysis indicates that during differentiation and maturation, astrocytes first downregulate programmes involved in proliferation and neuronal and glial development, before inducing genes involved in astrocyte functions in the adult brain. The induction of mature striatal astrocyte genes begins at early postnatal stages and continues into adulthood.

### Comparison of striatal and cortical datasets reveals a common astrocyte maturation signature

We next performed a bulk RNA-Seq analysis of astrocytes isolated from the grey matter of the cerebral cortex at P4 when astrocytes are still immature, and at 2 months of age when they are mature (Figure 2A). We performed this complementary analysis using cortical astrocytes to determine which maturation-regulated genes identified by the scRNA-Seq approach are part of a generic astrocyte maturation programme and to exclude genes with a striatum-restricted expression.

**Figure 2:**
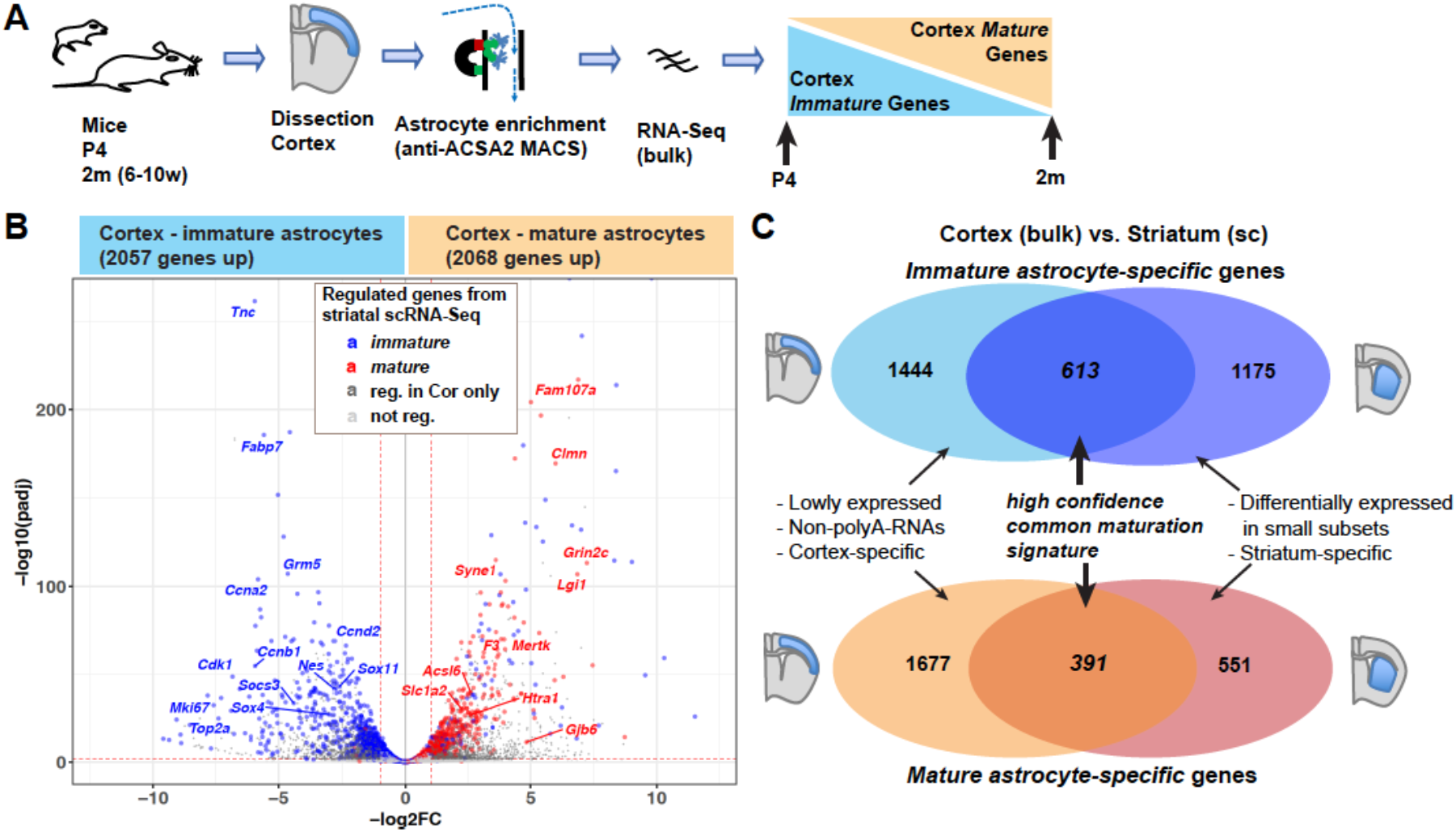
Comparison of striatal and cortical datasets reveals a common astrocyte maturation signature. **(A)** Approach used for MACS enrichment and bulk RNA-Seq analysis of immature astrocytes from P4 cortex and mature astrocytes from adult cortex. **(B, C)** Vulcano plot (B) and Venn diagrams (C) show genes differentially expressed between cortical astrocyte preparations from P4 and adult brain. Maturation-regulated genes identified by scRNA-Seq of striatal astrocytes (see Figure 1F) are highlighted in (B). Venn diagrams (C) show overlap of genes significantly regulated between early postnatal and adult astrocyte preparations in both striatal scRNA-Seq and cortical RNA-Seq datasets. Biological and technical differences between the analyses may contribute to the differences in genes detected in the two datasets, as indicated in (C). Differential genes: DESeq2 analysis (each n=3; padj ≤ 0.05, abs(log2FC) ≥1). See also Table S2.

Using DESeq2, we detected 2057 genes that were downregulated and 2068 genes that were upregulated in cortical astrocytes between P4 to 2 months (Fig. 2B/C, Table S2). These included many transcription factors and other genes expressed at low levels that were not detected in the scRNA-Seq analysis (Figure 2B/C, Table S2). Of the immature striatal astrocyte genes detected by scRNA-Seq analysis, 34% were significantly downregulated in cortical astrocytes between P4 and 2 months. Reciprocally, 42% of the mature striatal astrocyte gene sets were significantly upregulated in adult cortical astrocytes, including well-characterised and functionally important astrocytic genes such as *Slc1a2*, *Mertk* and *Gjb6*. Most other immature and mature striatal astrocyte genes showed a similar trend of regulation during maturation of cortical astrocytes, although not reaching statistical thresholds (Figure 2B/C, Table S2).

This analysis suggests that large parts of the astrocyte maturation gene expression programme identified by scRNA-Seq in the striatum also operate during maturation of cortical astrocytes. The 391 genes that were upregulated between early postnatal astrocytes and adult astrocytes in both brain regions irrespective of the different designs of the two experiments represent a high confidence set of *mature astrocyte-specific genes* (from hereon also referred to as *mature genes* for brevity). Similarly, the 613 genes that were consistently downregulated represent high confidence *immature astrocyte-specific genes* (from hereon also called *immature genes*).

### Maturation-associated chromatin remodelling at putative regulatory elements

To gain insights into the molecular control of astrocyte maturation, we next investigated the mechanisms that may regulate the differential expression of the *immature astrocyte-specific* and *mature astrocyte-specific* sets of genes. Chromatin remodelling establishes stable transcriptional changes during cellular differentiation processes. In particular, chromatin opening is required for non-pioneer transcription factors to bind and activate cell-type-specific enhancers (Perino and Veenstra, 2016). We investigated the changes in chromatin accessibility taking place during astrocyte maturation by performing *assay for transposase-accessible chromatin using sequencing* (Buenrostro et al., 2013) in cortical astrocytes isolated at P4 and 2 months (Figure 3A). A total of 95,066 peaks were called on the merged ATAC-Seq dataset. Removal of small peaks that were not clearly distinguishable from the background retained a total of 56,219 peaks (see methods and Figure S2A). 56% of the astrocyte ATAC peaks overlapped with reference DNase-Seq peaks from brain tissue at embryonic day 18.5 and in adult mice (Encode database, https://www.encodeproject.org; Figure S2B/C), as an alternative method to detect open chromatin, demonstrating the high quality of the data. Many ATAC peaks covered annotated transcriptional start sites (TSS) (called below “promoters”) but most were located in intergenic regions and introns. 44% of TSS-distal ATAC peaks overlapped with the enhancer-enriched histone mark H3K4me1 in reference brain ChIP-Seq datasets (Encode database; Figure S2B/C), suggesting that a large fraction of distal ATAC peaks are located in enhancer elements (called below “enhancers”). To identify genes that might be regulated by these candidate enhancers, we used a closest TSS-based approach (Figure 3B, S3D/E). We also integrated published regulatory regions from mouse brain tissue (Ron et al., 2017; Shen et al., 2012), which identified additional potential enhancers located at great distance from the genes they regulate, e.g. distal enhancers of the *immature astrocyte-specific* gene *Sox4* (Figure 3B, S3D/E, Table S3). Using the ATAC-Seq data, annotations in the Ensembl database and reference datasets, we compiled a list of 15,216 accessible promoters, 22,501 accessible enhancers and 18,502 additional distal accessible regions in both immature and mature astrocytes (Figure S2F, Table S3).

**Fig. 3:**
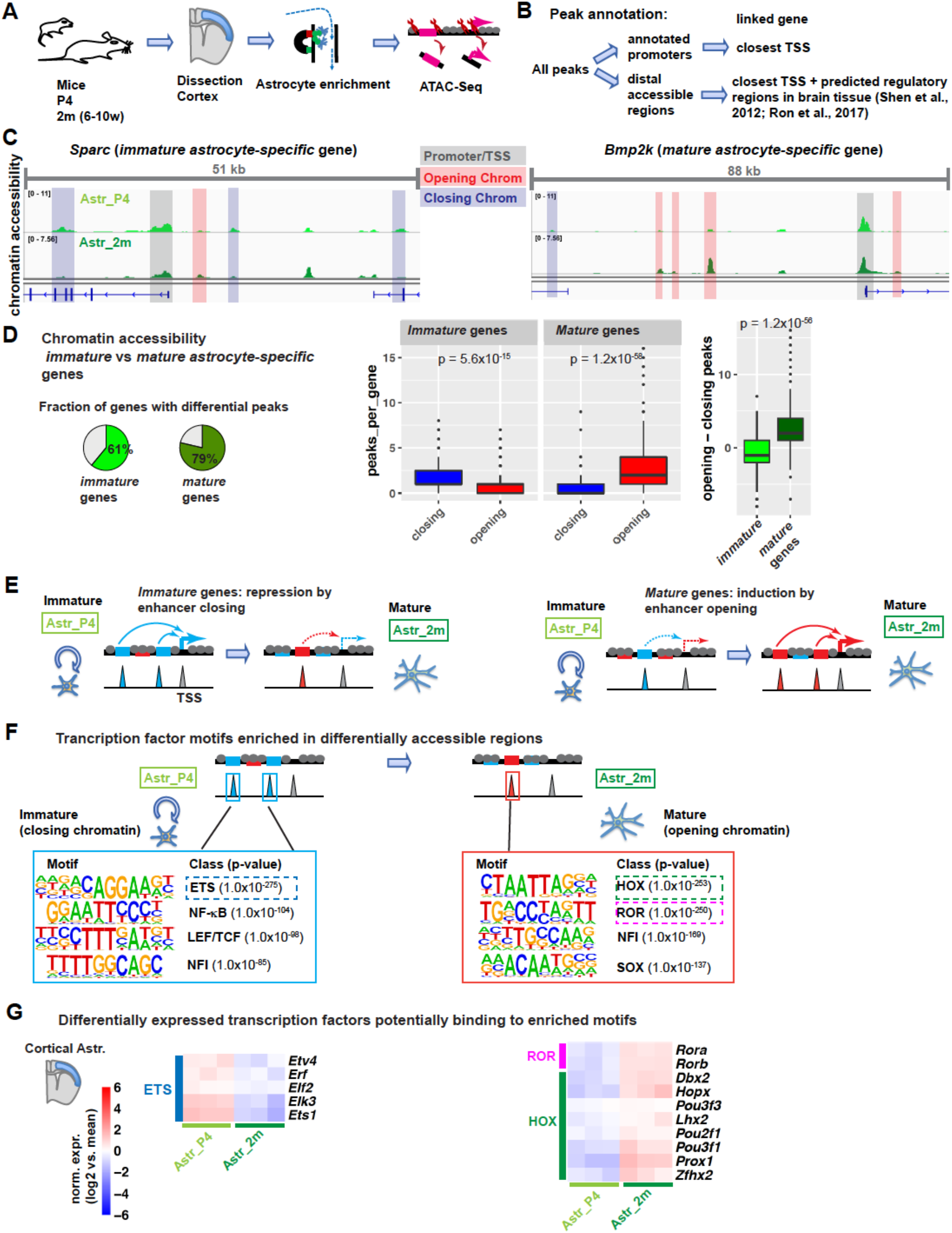
Maturation-associated chromatin remodelling at putative regulatory elements. **(A)** Approach used for MACS enrichment and ATAC-Seq analysis of immature and mature astrocytes from P4 and 2-month-old cortices, respectively (Astr_P4, Astr_2m), in order to identify stage-specific accessible chromatin regions, i.e. putative regulatory elements. **(B)** Strategy for mapping ATAC-Seq peaks to putative target genes. **(C)** Genome tracks show chromatin accessibility around the transcription start sites (TSS) of the *immature astrocyte-specific* gene *Sparc* and of the *mature astrocyte-specific* gene *Bmp2k*. Reduced size of ATAC-Seq peaks in adult vs P4 astrocytes indicates closing chromatin, while increased peak size indicates opening chromatin. **(D, E)** Genes regulated during astrocyte maturation (from Figure 2C) are globally associated with chromatin regions that change in their accessibility between P4 and adult stages (D). This suggests a model whereby changes in enhancer accessibility contribute to changes in gene expression during astrocyte maturation (E). **(F)** Transcription factor binding motifs enriched in closing and opening chromatin, identified by de novo motif enrichment analysis. The most enriched motifs are similar to ETS, homeodomain (HOX) and retinoic acid receptor related orphan receptor (ROR) transcription factor binding motifs. **(G)** Expression in cortical astrocytes of selected transcription factors that may bind to motifs highlighted in Fig. 3F based on bulk RNA-Seq analysis in Figure 2. The factors displayed are differentially expressed either in the cortical (bulk) or the striatal (single cell) RNA-Seq dataset and show a similar trend in the other dataset. Expression of the factors in striatal astrocytes is shown in Figure S2G. Statistical analysis and data presentation: (C) Tracks represent merged reads of 3-4 replicates; (C, D) Differential peaks: DESeq2 analysis (padj ≤ 0.05, abs(log2FC) ≥1); (D) Wilcoxon Rank Sum test (n= 355 immature vs 291 mature genes with associated differential peaks). See also Figure S2, Tables S3, S4, Supplementary Data S1.

Comparing the ATAC-Seq data from immature and mature astrocytes revealed prominent changes in chromatin accessibility between these stages, particularly in distal accessible regions (Figure 3C, S2D/F). Differentially accessible regions, determined by comparing ATAC-Seq read counts per peak using DESeq2, were associated with 61% of *immature astrocyte-specific* genes, such as *Sparc*, and 79% of *mature* genes, such as *Bmp2k* (Figure 3C/D, Table S4). *Immature* genes were predominantly associated with regions less accessible at 2 months than at P4 (closing chromatin) while *mature* genes were predominantly associated with elements more accessible at 2 months (opening chromatin; Figure 3C/D). Downregulated (*immature*) genes had a median of 1 more associated ATAC peak closing than opening, while upregulated (*mature*) genes had a median of 2 more peaks opening than closing (p = 1.2×10^−56^, Wilcoxon test; Figure 3D, Table S4). These results suggest that changes in enhancer accessibility may contribute to the transcriptional changes taking place during astrocyte maturation (Figure 3E).

We next aimed to identify transcription factors that may drive changes in gene expression underlying astrocyte maturation. For this, we performed motif enrichment analysis in differentially accessible regions using the Homer software. In chromatin regions that closed between P4 and 2 months, the most enriched binding motifs included those for ETS, NF-κB and LEF/TCF transcription factors (Figure 3F). In chromatin regions that opened between postnatal and adult stages, motifs associated with homeodomain (HOX) and retinoic acid receptor-related orphan receptor (ROR) factors were most enriched (Figure 3F). Motifs for the key astrogliogenic NFI transcription factors were found enriched in both opening and closing chromatin (Figure 3F).

Examination of the cortical bulk RNA-Seq dataset (Figure 3G) and/or the striatal sc-RNA-Seq dataset (Figure S2G) for the expression of transcription factors associated with the enriched motifs, revealed that several ETS genes had a lower expression in adult than early postnatal astrocytes. Reciprocally, *Rora*, *Rorb*, and several HOX genes including *Dbx2*, *Lhx2* and *Pou3f3* (*Brn1*) were upregulated in adult astrocytes (Figure 3G, S2G). Altogether, our data suggest that transcription factors from the ETS, HOX and ROR families may direct chromatin remodelling events and thereby contribute to the transcriptional changes associated with astrocyte maturation.

### *In vitro* differentiation of NSCs into astrocytes fails to recapitulate *in vivo* maturation

To examine the activity of candidate transcription factors in promoting astrocyte maturation, we turned to an *in vitro* model of astrocyte differentiation. We established NSC cultures from the early postnatal cortex, a likely source of cortical astrocytes *in vivo* (Ge et al., 2012), expanded them in the presence of EGF and FGF2, and subsequently differentiated them into astrocytes by growth factor withdrawal and addition of BMP4 (Bonaguidi et al., 2005; Kleiderman et al., 2015) (Figure 4A).

**Fig. 4:**
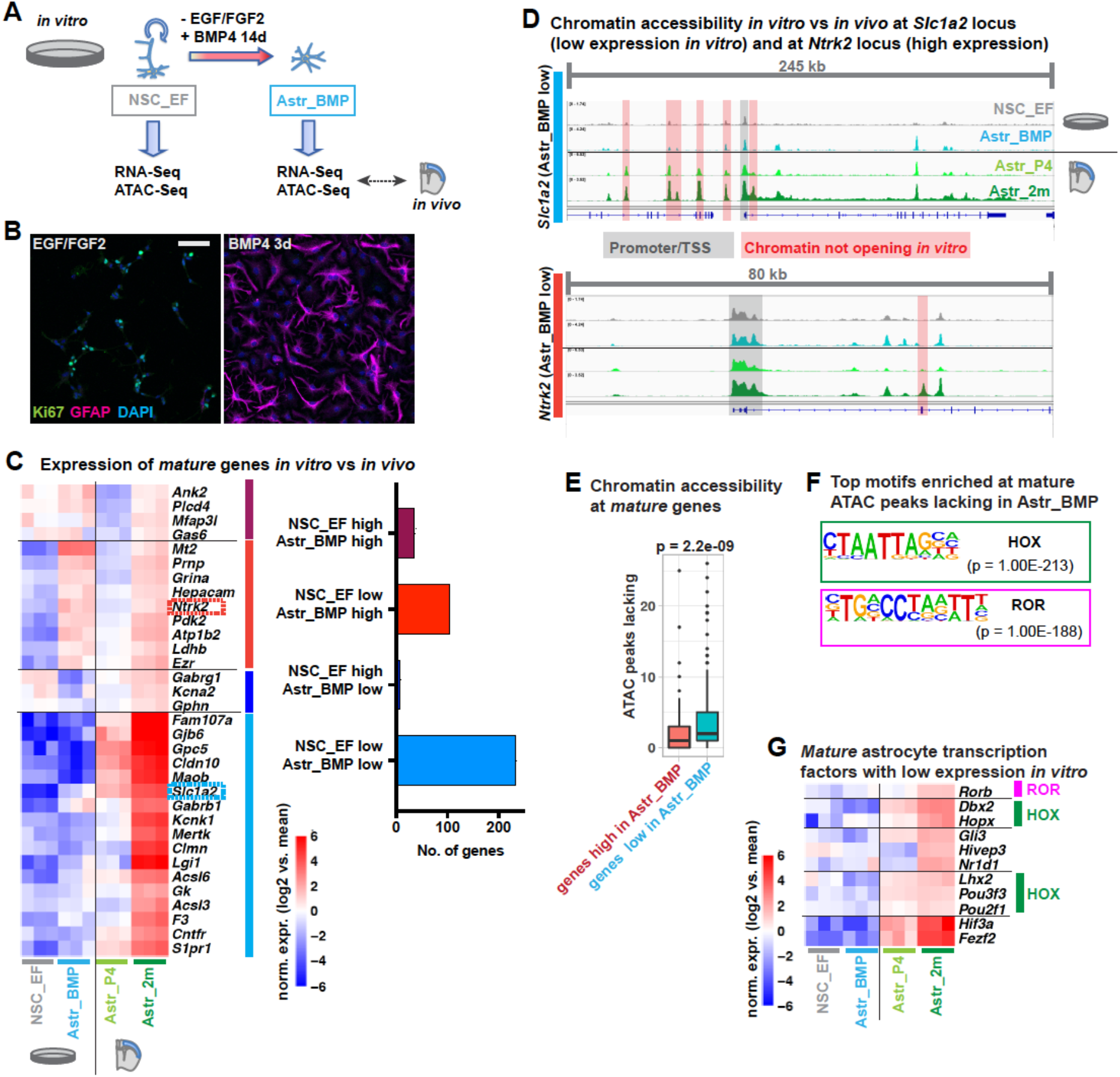
*In vitro* differentiation of NSCs into astrocytes fails to recapitulate *in vivo* maturation. **(A)** NSCs maintained in EGF/FGF2-containing medium (NSC_EF) are differentiated into astrocytes by culture in BMP4-containing medium for 14 days (Astr_BMP); maturity of astrocytes in these cultures is assessed by RNA-Seq and ATAC-Seq analysis and comparison with datasets from P4 and adult cortical astrocytes (Astr_P4/Astr_2m from Figures 2 and 3). **(B)** Immunolabeling for the proliferation marker Ki67 and the astrocyte marker GFAP after 3 days of differentiation; scale bar 50 um. **(C)** Expression of *mature astrocyte-specific* genes (from Figure 2C) in cultured astrocytes and brain astrocytes. Heatmap of selected genes and number of genes, grouped by their expression patterns in cultured astrocytes. “NSC_EF/Astr_BMP low” refers to significantly lower expression compared to Astr_2m, “high” to equal or higher expression. **(D)** ATAC-Seq genome tracks show chromatin accessibility in cultured astrocytes vs brain astrocytes around the TSS of *Slc1a2* and *Ntrk2* (expression highlighted in (C)). **(E)** *Mature astrocyte-specific* genes with low expression *in vitro* (Astr_BMP) lack accessibility at sites open *in vivo* (Astr_2m), more than genes with high expression. **(F)** Transcription factor binding motifs enriched in DNA regions where ATAC-Seq analysis shows lower chromatin accessibility in cultured astrocytes compared to adult brain astrocytes (de novo motif enrichment analysis). **(G)** Expression of selected transcription factors induced during astrocyte maturation in vivo and expressed at low levels in cultured astrocytes. Highlighted are factors that might bind the motifs in (F). Heatmap of cortical astrocyte bulk RNA-Seq data from Figure 2; the factors displayed are differentially expressed in the cortical (bulk) or striatal (single cell) RNA-Seq dataset and show a similar trend in the other dataset. Expression of the factors in striatal astrocytes is shown in Figure S3D. Statistical analysis and data presentation: Differential genes/peaks: padj ≤ 0.05, abs(log2FC) ≥ 1, (DESeq2 analysis); (D) Genome tracks of 3-4 merged replicates. (E) Wilcoxon Rank Sum test (n=233 with low vs 138 genes with high expression *in vitro*). See also Figure S3, Table S5.

After 3 days in differentiation medium, NSCs had downregulated the proliferation marker Ki67 and begun to express the astroglial lineage marker GFAP (Figure 4B). Cortex-derived NSCs, as well as adult subventricular zone NSCs, a well characterised reference population, were differentiated into astrocytes in the presence of BMP4 for 14 days to allow time for maturation, and were then analysed by RNA-Seq. Their transcriptional profile was compared with that of undifferentiated NSCs in culture and with the RNA-Seq datasets obtained from cortical astrocytes acutely isolated from mice at P4 and 2 months (Figure 2).

Many astrocyte markers were upregulated in *in vitro*-differentiated astrocytes compared to NSCs, including *Aldh1l1*, *Sox9*, *Gfap*, and *Aqp4*, which reached expression levels comparable to those observed in cortical astrocytes in 2-months old brains (Figure S3A). To compare the maturation state of *in vitro*-differentiated astrocytes with that of adult brain astrocytes, we next analysed the expression of *immature* and *mature astrocyte-specific* genes (Figure 2). Astrocytes differentiated *in vitro* expressed 37% of *mature* genes at levels similar to those observed at 2 months *in vivo*, but 63% were expressed at lower levels, including key functional effector genes such as *Gjb6*/*Cx30* and *Slc1a2*/*GLT-1* (Figure 4C, Table S5).

Compared to the limited induction of *mature* genes, the differentiation protocol was more efficient at repressing *immature* genes. *In vitro*-differentiated astrocytes expressed 75% of *immature* genes at low levels, similar to those found in *mature* astrocytes *in vivo* and only 25% were expressed at high levels (Fig. S3B). Comparison of our dataset for *in vitro*-differentiated astrocytes with published data for cultured primary astrocytes (Hasel et al., 2017) and astrocytes differentiated with CNTF (Tiwari et al., 2018) showed that astrocytes in different *in vitro* models present similarly low expression levels of many *mature astrocyte-specific* genes (Fig. S3C). Overall these results suggest that *in vitro*-differentiated astrocytes do not reach a level of maturation comparable to that of adult brain astrocytes.

We next performed a comparative ATAC-Seq analysis of *in vitro*-differentiated astrocytes and NSCs to investigate changes in chromatin accessibility associated with *in vitro* differentiation and compare accessible sites with those observed in adult brain astrocytes. At the *Slc1a2* locus, a *mature* gene expressed only at low levels *in vitro* (Figure 4C/D), many sites which had large ATAC peaks in adult brain astrocytes presented only small signals in cultured astrocytes (Fig. 4D). In contrast, the *Ntrk2* locus, which is expressed highly in both *in vitro*-differentiated and adult brain astrocytes, showed more similar accessibility profiles in the *in vitro* and brain samples (Figure 4C/D). Globally, *mature* genes that were expressed at low levels in *in vitro*-differentiated astrocytes were lacking more accessibility peaks than genes highly expressed in cultured cells (Fig. 4E). Motif enrichment analysis revealed that the sequences most enriched in regions with reduced accessibility in cultured astrocytes were HOX- and ROR-binding motifs (Fig. 4F), which were also enriched in chromatin regions opening in brain astrocytes between P4 and 2m (Fig. 3F). Together our results show that *in vitro*-differentiated astrocytes fail to gain chromatin accessibility at many regulatory elements associated with *mature astrocyte-specific* genes. The results also suggest that a lack of activity of ROR and HOX transcription factors in *in vitro*-differentiated astrocytes might prevent chromatin opening at key regulatory regions and may block full maturation. Supporting this hypothesis, we found that several ROR and HOX genes induced during maturation *in vivo* remained expressed at low levels in *in vitro*-differentiated astrocytes, including *Rorb*, *Dbx2* and *Lhx2* (Figure 4G, S3D).

### The astrocyte maturation programme is comprised of several transcriptional modules

We next asked whether individual transcription factors that are induced during astrocyte maturation regulate the *mature* astrocyte phenotype, and whether the failure of some of these factors to be expressed in cultured astrocytes contributes to the lack of maturation of these cells. To address these questions, we performed a reconstitution assay in *in vitro*-differentiated astrocytes, whereby we forced expression of candidate transcription factors that normally remain expressed at low levels in these cells. We then determined whether expression of each of these factors was sufficient to induce parts of the maturation programme that fail to be induced during *in vitro* differentiation. We focussed on four candidates, the ROR protein Rorb, the HOX proteins Dbx2 and Lhx2 and the zinc finger protein Fezf2, which do not reach mature *in vivo* levels during *in vitro* differentiation (Figure 4G).

We expressed the candidate regulators in NSC-derived astrocytic cultures after 7 days in differentiation medium using a lentiviral expression system (Figure 5A). Efficiency of the delivery method was demonstrated by immunolabeling for the the V5-tagged factors and for the mature astrocyte protein glutamine synthetase (GS/Glul), which was induced by both *Rorb* and *Fezf2* (Figure 5B). To investigate the transcriptional activity of the candidate regulators, we performed an RNA-Seq analysis of control and transcription factor-transduced *in vitro*-differentiated astrocytes, and compared these transcript profiles with those of postnatal and adult cortical astrocytes (Figure 5C, S4A-B, Table S6). Both *Rorb* and *Fezf2* induced and repressed several hundred genes, whereas *Dbx2* and *Lhx2* regulated smaller gene sets and predominantly activated transcription (Figure S4A). Gene Ontology analysis revealed that the gene sets induced by the transcription factors were all enriched for the terms “blood circulation” and “calcium homeostasis” and related processes linked to astrocyte function. GO terms associated with *Rorb*- and *Lhx2*-induced genes also included “regulation of synaptic plasticity” and “axonogenesis” and *Fezf2*-induced genes were associated with “learning or memory” and “regulation of neurotransmitter levels” (Figure S4A, Table S6). Many of the transcription factor-induced genes were among the *mature astrocyte-specific* genes expressed only at low levels in control cultures (Figure 5C). *Rorb* alone induced 14% of the *mature* genes (36 out of 253 genes), including genes with well-established roles in astrocyte physiology such as *Glul* and *Apoe*, while *Fezf2* induced 10% (25 genes), *Dbx2* 4% (10 genes) and *Lhx2* 4% (9 genes) of this gene set (Figure 5C). Interestingly, the genes regulated by the different transcription factors were largely non-overlapping, although they were often linked to related astrocyte functions. For example, *Apoe* (induced by *Rorb*), *S100b* (induced by *Fezf2*), and *Atp1a2* (induced by *Dbx2* and *Fezf2*) have all been reported to regulate astroglial calcium signalling (Golovina et al., 2003; Muller et al., 1998; Xiong et al., 2000). Similarly, *Atp1a2* and the *Lhx2*-induced adenosine receptor *Adora2b* have both been implicated in the regulation of vasoconstriction (Busse et al., 2016; Hermann et al., 2013).

**Figure 5:**
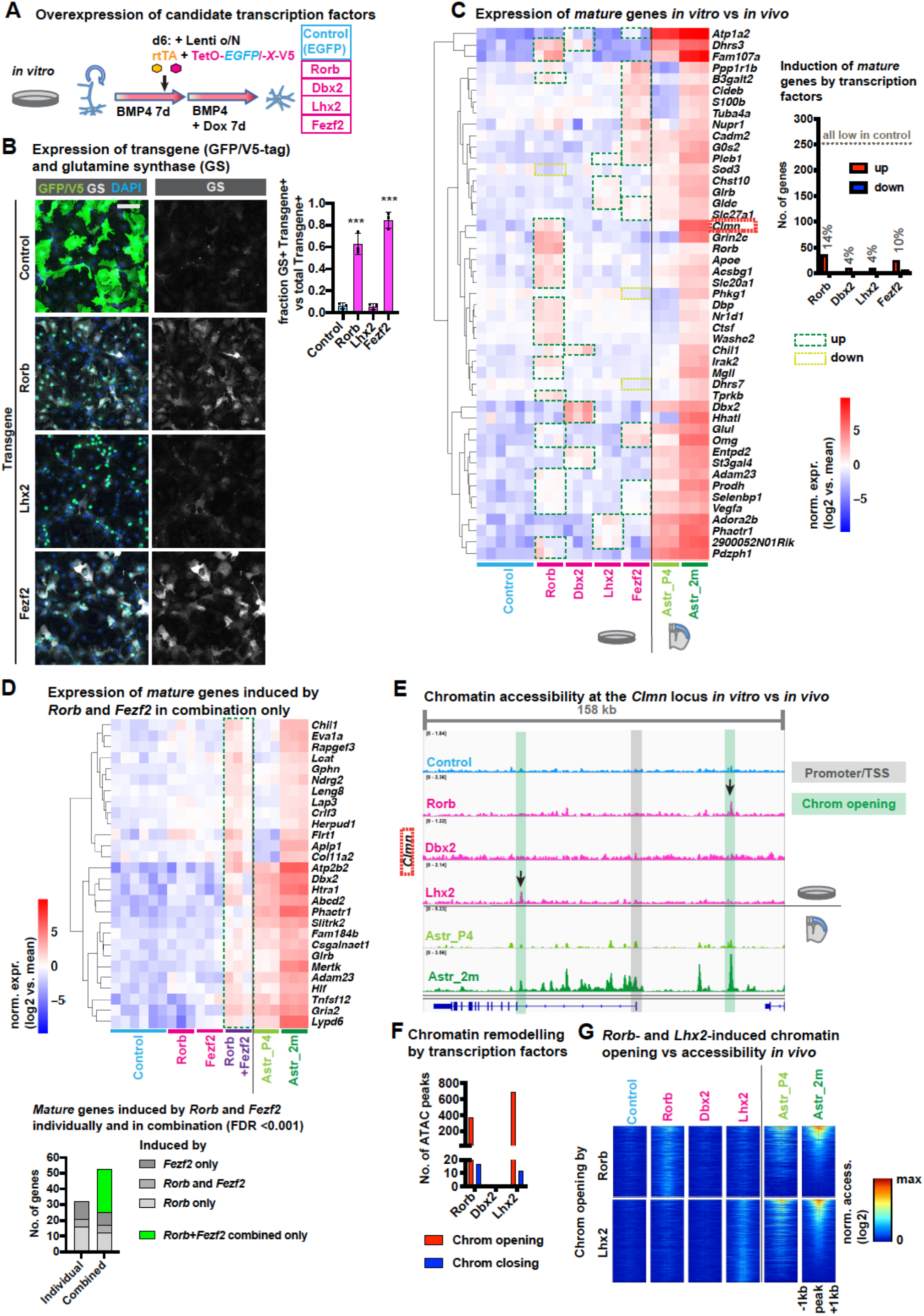
The astrocyte maturation programme is comprised of several transcriptional modules. **(A)** Doxycycline-inducible expression of V5-tagged transcription factors (TFs) or EGFP as a control, in astrocyte cultures to assess the contribution of *Rorb*, *Dbx2*, *Lhx2* and *Fezf2* to astrocyte maturation. Cells are infected with the lentiviral vectors at day 6 of differentiation, followed by transgene induction with doxycycline from day 7 to day 14. **(B)** Immunolabeling for the transgene (EGFP or V5 tag) and the mature astrocyte enzyme glutamine synthase (GS, gene Glul). **(C)** Expression of *mature astrocyte-specific* genes (from Figure 2C) in cultured astrocytes and cortical astrocytes. Heatmap of selected genes. Barplot showing that subsets of *mature* genes with low expression in control cultures are induced by the transcription factors analysed. **(D)** Regulation of *mature astrocyte-specific* genes by the combined expression of *Rorb* and *Fezf2*. Astrocytes co-infected with *Rorb* and *Fezf2* viruses or with EGFP viruses were analysed along with samples from (C). **(E)** ATAC-Seq genome tracks showing chromatin accessibility peaks around the TSS of the *mature* gene *Clmn* in cultured astrocytes expressing transcription factors and in brain astrocytes. Note the highlighted peaks that are present in cultured astrocytes only when *Rorb* or *Lhx2* are expressed. **(F)** Number of ATAC-Seq peaks induced or suppressed by expression of transcription factors in cultured astrocytes. **(G)** Heatmaps showing chromatin accessibility in cultured astrocytes around *Rorb*- and *Lhx2*-induced peaks and in brain astrocytes. Statistical analysis and data presentation: (B) scale bar 50 μm; n=3 biological replicates, ***p < 0.001 vs EGFP (one-way ANOVA with Tukey’s multiple comparisons test). (C-G) Differential genes/peaks: abs(log2FC) ≥1, padj ≤ 0.05 (padj ≤ 0.001 in (D)) (DESeq2 analysis); (E) Genome tracks of 3-4 merged replicates. (F) Heatmaps show normalised ATAC read count of merged samples in 20bp windows +/− 1kb of peak centres. See also Figure S4, Table S6.

Since developmental genes are often regulated by multiple transcription factors acting cooperatively (Long et al., 2016), we examined a possible synergy between *Rorb* and *Fezf2* by co-expressing the two genes in *in vitro*-differentiated astrocytes. RNA-Seq analysis revealed that co-expression of *Rorb* and *Fezf2* induced a number of additional *mature* genes that were not induced by either factor individually (Figure 5D). When setting a low false discovery rate (FDR) threshold of 0.001, only 32 *mature* genes were induced by *Rorb* or *Fezf2* separately (16 by *Rorb*, 11 by *Fezf2*, and 5 by both). The two factors together induced almost twice this number (53 genes; Figure 5D). In addition to inducing *mature* genes, *Rorb* and *Fezf2* also repressed several *immature* genes whose expression was maintained in *in vitro*- differentiated astrocytes (Figure S4B). In combination, *Rorb* and *Fezf2* repressed additional *immature* genes, including progenitor markers and genes supporting neuronal development, such as *Nes*, *Vim*, *Fabp7*, *Sparc*, *Marcks* and *Marcksl* (Fig. S4C).

To address whether the regulation of gene expression by ROR and HOX proteins in *in vitro*-differentiating astrocytes involves changes in chromatin accessibility, we performed an ATAC-Seq analysis of cultured astrocytes expressing *Rorb*, *Dbx2* or *Lhx2*. Expression of *Rorb* and *Lhx2*, but not *Dbx2*, resulted in chromatin opening at the locus of the *mature astrocyte-specific* gene *Clmn* and several hundred other loci (Fig. 5E/F). Comparison of these ATAC-Seq profiles with those of adult cortical astrocytes *in vivo* showed that many sites that acquired an ATAC-Seq peak upon transcription factor expression *in vitro* also had an ATAC-Seq peak in adult astrocytes *in vivo* (Fig. 5E/G). Overall, our data strongly suggest that *Rorb*, *Dbx2*, *Lhx2* and *Fezf2*, whose expression is induced when astrocytes mature *in vivo*, significantly contribute to the transcriptional and epigenetic maturation of astrocytes.

### Extrinsic signals promote the transcriptional and epigenetic maturation of astrocytes

The lack of expression of *Rorb*, *Dbx2*, *Lhx2* and *Fezf2* in *in vitro*-differentiated astrocytes suggested that these cultures were lacking a signaling environment that permits expression of these *mature astrocyte-specific* transcription factors. Previous studies have reported that cell-cell-contacts (Li et al., 2019) and FGF signalling (Roybon et al., 2013) could improve the functional maturation of cultured astrocytes. We therefore asked whether these conditions might stimulate astrocyte maturity by inducing the expression of *Rorb*, *Dbx2*, *Lhx2* or *Fezf2*. For this, we differentiated astrocytes for 7 days with BMP4 and cultured the cells for 7 additional days in control conditions (basal medium) or in the presence FGF2. We also differentiated astrocytes in basal medium or in the presence of FGF2 in a three-dimensional (3D) environment to facilitate cell-cell-interactions (Figure 6A, see methods). RNA-Seq analysis of cultures in these different conditions revealed a significant increase in the expression of several *mature astrocyte-specific* transcription factors that remained expressed at low levels in control astrocyte cultures (Figure 6B). In particular, FGF2 in two-dimensional (2D) cultures induced *Dbx2* expression, 3D cultures induced *Rorb* and *Hopx* expression, while combining FGF2 and 3D conditions had the strongest effect and markedly induced the expression of *Rorb*, *Dbx2*, *Lhx2* and other factors (Figure 6B). These conditions and in particular FGF2 in 3D cultures also increased the expression of a large number of *mature astrocyte-specific* genes, including many important astrocyte effector genes such as *Slc1a2*, *Glul*, *Acsl6*, *Gja1*/*Cx43* and *Mertk* (Figure 6C, Table S7). ATAC-Seq analysis showed that sites in the *Slc1a2* locus that were accessible in adult cortical astrocytes but not in control cultures, became accessible in 3D cultures, particularly in the presence of FGF2 (Figure 6D). On a global level, we observed that the combination of FGF2 and a 3D environment induced chromatin accessibility in cultured astrocytes at sites that also become accessible when astrocytes become mature in the brain (Figure 6E). Together these findings suggest that ameliorating the signalling environment of *in vitro*-differentiating astrocytes can stimulate the expression of key transcriptional regulators which in turn promote astrocytic maturation, at least in part via chromatin remodelling at *mature astrocyte-specific* loci.

**Fig. 6:**
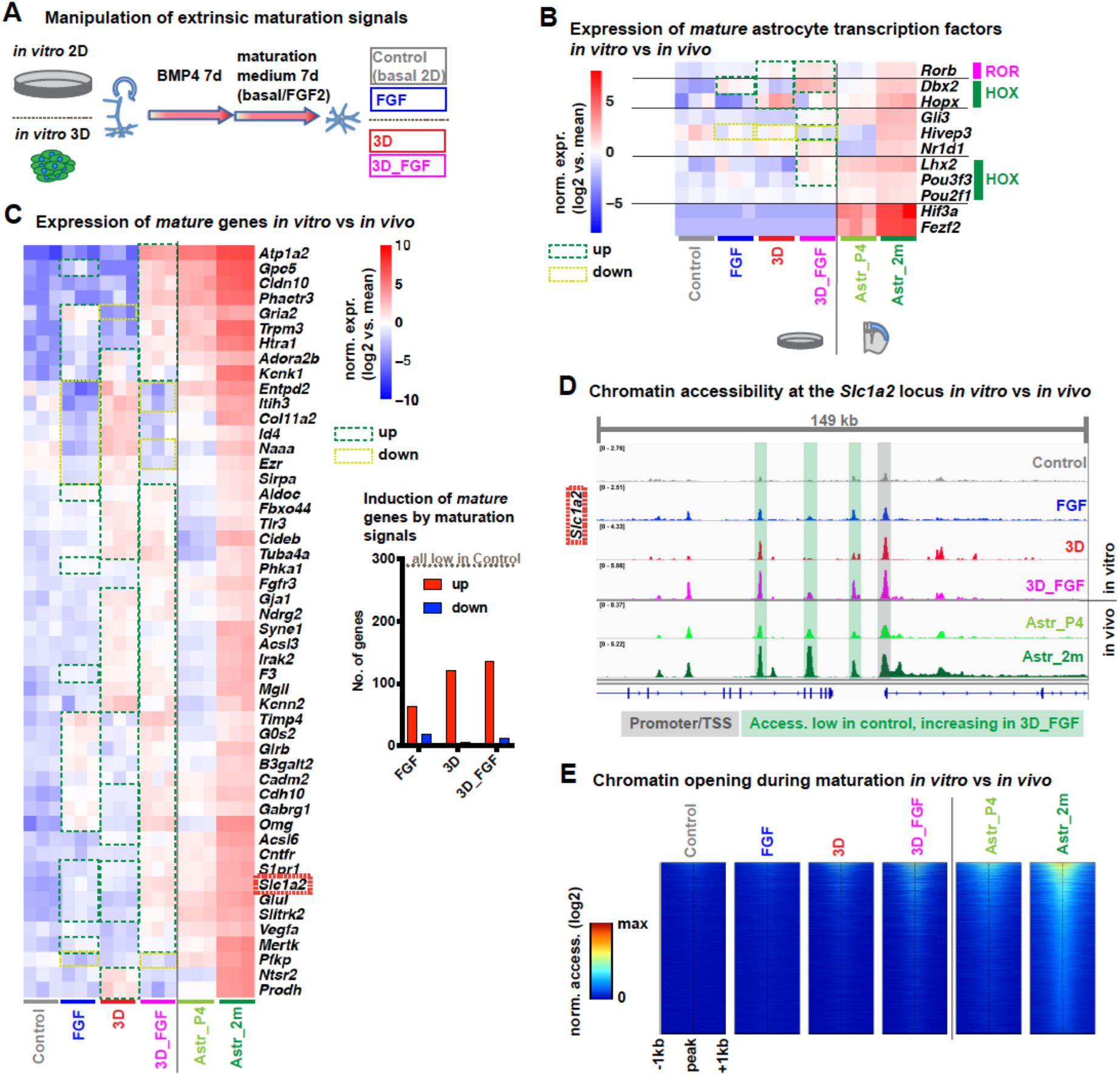
Extrinsic signals promote the transcriptional and epigenetic maturation of astrocytes. **(A)** Experimental design to assess the effects of extrinsic signals on astrocyte maturation. Initial differentiation with BMP4 is followed by 7 days in different maturation conditions: control (basal medium in conventional two-dimensional cultures) +/− FGF2 (control/FGF) or in three-dimensional gel-embedded cultures (3D/3D_FGF). **(B-C)** Expression of *mature astrocyte-specific* transcription factors (B) and other genes (C) in cultured astrocytes and cortical astrocytes. Heatmaps of selected genes. Barplot showing that subsets of *mature* genes with low expression in control cultures are induced in other culture conditions. **(D)** ATAC-Seq genome tracks showing chromatin accessibility peaks around the TSS of the *mature astrocyte-specific* gene *Slc1a2*, which is expressed at higher levels in astrocytes in 3D cultures in the presence of FGF2 than in astrocytes in control cultures (see (C)). **(E)** Heatmaps showing global chromatin accessibility around ATAC peaks that are induced during maturation of cortical astrocytes and in astrocytes in 3D cultures with FGF2, but are absent in control cultures (5499 peaks). Statistical analysis and data presentation: Differential genes/peaks: abs(log2FC) ≥ 1, padj ≤ 0.05 (DESeq2 analysis); (D) Genome tracks of 3-4 merged replicates. (E) Heatmaps show normalised ATAC read count of merged samples in 20bp windows +/− 1kb of peak centres. See also Table S7.

## Discussion

Astrocyte maturation, the last stage of astrocyte differentiation, takes place from early postnatal to young adult stages, when proliferating immature astrocytes cease to divide and aquire the full spectrum of astrocyte functions observed in the adult brain (Molofsky and Deneen, 2015; Yang et al., 2013). In this study, we have analysed in detail the transcriptional and chromatin changes that occur during maturation of astrocytes in the mouse forebrain. We found that defined extracellular signals can induce in cultured astrocytes the expression of transcription factors associated with astrocytic maturation, which are otherwise lacking. These transcription factors in turn induce chromatin remodelling and the expression of large numbers of genes associated with astrocyte maturity, which are organised in distinct regulatory modules activated by different transcription factors acting alone or in combination.

The global changes in gene expression occurring during astrocyte maturation have begun to be investigated only recently (Sloan et al., 2017; Tiwari et al., 2018; Zhang et al., 2016), and their interpretation is complicated by two limitations: first, because of the phenotypic similarity of neural stem cells and astrocytes, it is difficult to experimentally isolate or even define cells that are considered immature astrocytes, as opposed to neural stem cells or glial progenitors. Second, two of the papers (Sloan et al., 2017; Tiwari et al., 2018) are based on *in vitro* models that do not fully recapitulate maturation *in vivo*. Nevertheless, these recent studies have revealed major transcriptional changes occurring during late stages of differentiation or maturation, in agreement with the findings in this study. However single cell RNA-Seq analysis allowed us to resolve gene expression changes with greater resolution than in previous studies. We propose in particular that immature proliferating astrocytes or astrocyte progenitors progress to a fully mature state through at least three intermediate stages in which astrocytes have exited the cell cycle but are still immature. It is currently unclear to what extent these intermediate transcriptional stages represent cells with different functional properties, e.g. supporting different steps of neuronal development.

Single cell transcriptome analysis showed that during astrocyte differentiation and maturation, *immature astrocyte-specific* genes linked to proliferation and neuronal and glial development were downregulated first, before *mature astrocyte-specific* genes were induced in two waves. The first wave of *mature* genes was induced in early postnatal astrocytes and they retained high expression levels in adult astrocytes. The second wave of *mature* genes reached highest expression levels only in adult astrocytes and included genes with important roles in astrocyte physiology such as *Slc1a2*/*GLT-1*, a glutamate transporter that prevents hyperexcitation of glutamatergic synapses, and *Gjb6*/*Cx30*, which is required for the removal of potassium and the rapid supply of metabolites to active neurons (Dallerac et al., 2018). Thus our analysis emphasises the importance of the step of maturation that takes place between P7 and 2 months of age, for the acquisition of the full spectrum of adult astrocyte functions.

To investigate the genomic mechanisms underlying astrocyte maturation, we identified putative regulatory elements present in immature or mature astrocytes *in vivo*, by mapping accessible chromatin regions with ATAC-Seq. Integrating these datasets with gene expression profiles suggested that chromatin remodelling has an important role in the transcriptional changes taking place during astrocyte maturation. Motif enrichment in elements that become inaccessible when astrocytes mature suggests that several families of transcription factors acting downstream of extracellular signals maintain immature astrocyte-specific enhancers in an open state and thus support the unique properties of immature astrocytes, including perhaps their capacity to reactivate NSC programmes and become proliferative and neurogenic. These factors include ETS factors, whose activity is regulated by receptor tyrosine kinase receptor signalling (Newton et al., 2018; Sizemore et al., 2017), the Wnt pathways effectors LEF/TCF (Cadigan and Waterman, 2012), and NF-κB acting downstream of inflammatory signals (Ben-Neriah and Karin, 2011; Lattke and Wirth, 2017). The limited availability of these developmental signals in the adult brain might contribute to the loss of lineage plasticity in mature astrocytes. In the future it will be interesting to determine whether reactivating these developmental signals or re-expressing the downstream transcription factors could restore plasticity to mature astrocytes.

Conversely, we found HOX and ROR transcription factor-binding motifs enriched in chromatin regions that become accessible during maturation. The expression of several members of these families also increased during maturation, including the ROR factor *Rorb*, and the HOX factors *Dbx2* and *Lhx2*, suggesting that induction of these transcription factors might result in the opening of enhancers required for the expression of *mature astrocyte* genes.

To functionally dissect the mechanisms underlying astrocyte maturation, we used an *in vitro* model of differentiation of cultured NSCs into astrocyte in the presence of BMP4 (Bonaguidi et al., 2005; Kleiderman et al., 2015). We found that astrocytes differentiated with this protocol, as well as astrocytes differentiated with CNTF (Tiwari et al., 2018) and cultured primary astrocytes (Hasel et al., 2017), remained partially immature. This finding highlights the difficulty of studying astrocyte maturation and physiology in culture, a serious handicap for this field (Hasel et al., 2017; Krencik et al., 2011; Roybon et al., 2013). This lack of maturation has recently been ascribed to the absence of relevant extrinsic signals, including growth factors and contacts with neurons and other astrocytes (Hasel et al., 2017; Li et al., 2019; Roybon et al., 2013). However a limitation of these studies is that the maturity of *in vitro*-differentiated astrocytes was not compared with that of mature brain astrocytes. Here we found that three-dimensional cultures, which allow more extensive cell-cell-contacts, showed improved astrocyte maturity, particularly in combination with FGF2. However, even in these conditions *in vitro*-differentiated astrocytes are not fully mature and still lack expression of many maturity-associated genes. These findings show that our approach of comparing the molecular profile of *in vitro*-differentiated astrocytes with that of astrocytes acutely isolated from the brain is a powerful strategy to further improve differentiation protocols for cultured astrocytes.

While the incomplete maturation of astrocytes *in vitro* limits their suitability as models to study astrocyte physiology and function, it provided an excellent assay to investigate the mechanisms that promote maturation. By reconstituting the expression of transcription factors that are induced in astrocytes maturing *in vivo* but are missing from cultured astrocytes, we could demonstrate a maturation-promoting activity of four factors, *Rorb*, *Dbx2*, *Lhx2* and *Fezf2*. These factors have mainly been studied for their roles in neuronal subtype specification (Chen et al., 2008; Muralidharan et al., 2017; Oishi et al., 2016; Pierani et al., 1999). Interestingly, their roles in neurons seem to be mediated at least partially by cross-repression of alternative neuronal subtype programmes, suggesting that their expression in astrocytes might contribute to the global suppression of neuronal programmes. Moreover *Dbx2* and *Fezf2* have also been shown to promote NSC quiescence (Berberoglu et al., 2014; Lupo et al., 2018), thus these two factors might contribute to suppressing the proliferative capacity of mature astrocytes. In addition to such repressive activities, each of the four factors also induced maturity-associated genes in cultured astrocytes, suggesting that they may also have transactivating functions in the astrocytic context.

Interestingly, the four factors induced largely non-overlapping sets of astrocyte effector genes. The induction of distinct sets of maturity-associated gene by individual transcription factors suggests that the astrocyte maturation programme is regulated in a modular manner. This is different from cell fate specification and early differentiation steps, where individual master regulators trigger a highly coordinated cell intrinsic differentiation programme. For example, MyoD is able to induce myogenic programmes in a variety of cell types (Weintraub et al., 1989), and NFIA can trigger astrocyte differentiation of human NSCs (Tchieu et al., 2019). In contrast, our results suggest that maturation is controlled by a large number of transcription factors which induce astrocytic effector subprogrammes by both independent and cooperative activities. This distinct organisation of the maturation programme may be explained by different regulatory requirements. Cell fate specification and early differentiation processes are defining the fundamental properties of a cell type largely independently of extrinsic conditions, and are therefore relying mostly on cell intrinsic master regulators and downstream transcription factor networks. In contrast, maturation programmes are involved in shaping the cellular effector functions and must therefore be closely adapted to the changing requirements of the organism, particularly for a highly plastic and multifunctional cell type such as astrocytes. The regulatory adaptility of a cell is best maintained if its different functions are controlled independently by different transcription factors that are responsive to distinct extrinsic factors that relay various physiological requirements. This modular maturation model is supported by our results showing that astrocytes respond to FGF2 and three-dimensional cultures by inducing the transcription factors *Rorb*, *Dbx2* and *Lhx2*, which in turn induce different sets of mature astrocyte genes. To what extent this model can be generalised to other cell types remains to be investigated, but a similar model has also been discussed for the differentiation and maturation of T lymphocyte subtypes, which are determined by a complex regulatory network involving cytokines as extrinsic signals and a spectrum of transcription factors rather than individual master regulators (Chang et al., 2014).

Importantly, we also found that the maturation-associated transcription factors do not act fully independent but cooperate on several levels. First, they induce distinct sets of effector genes that may regulate the same astrocyte functions. For example, *Apoe* (induced by *Rorb*), *S100b* (induced by *Fezf2*), and *Atp1a2* (induced by *Dbx2* and *Fezf2*) have all been reported to regulate intracellular calcium levels in astrocytes (Golovina et al., 2003; Muller et al., 1998; Xiong et al., 2000). Second, the induction of chromatin remodelling by *Rorb* and *Lhx2* might be a prerequisite for other factors to promote the expression of astrocytic effector genes. In particular *Lhx2* induces chromatin opening at a large number of sites but regulates only few genes, suggesting that *Lhx2*-dependent chromatin opening may “bookmark” genes for future induction by other factors, as has been recently reported for *C/EBPa* and *HNF4a* in liver development (Karagianni et al., 2020). Such a mechanism might contribute to the observed cooperativity of *Rorb* and *Fezf2*, which together induced the expression of many more genes than were induced by each factor separately. One potential explanation for this cooperative activity might be that *Rorb* opens enhancers that are subsequently bound and activated by *Fezf2*. Alternatively, the two factors might bind simultaneously to the same or different enhancers of the same gene (Long et al., 2016).

Additional research is required to fully determine the functions of the transcription factors investigated here, in promoting specific aspects of the mature phenotype of astrocytes. *Lhx2* has been shown to contribute to the differentiation of tanycytes and the maintenance of the mature state of Müller glia, two astrocyte-like cell types found in the hypothalamus and the retina, respectively (de Melo et al., 2012; Salvatierra et al., 2014), but *Rorb*, *Dbx2* and *Fezf2* have not been previously implicated in the development of glial cells. Interestingly, pathogenic mutations in *RORB* have been identified in genetic forms of epilepsy (Rudolf et al., 2016). Since astrocyte dysfunction is considered a key pathological mechanism in epilepsy (Patel et al., 2019), these *RORB* mutations may act by blocking the induction of mature astrocyte genes protecting from epileptogenesis (Patel et al., 2019), such as the *Rorb* target *Glul*, whose deletion in astrocytes in the mouse causes seizures and neurodegeneration (Zhou et al., 2019), or the glutamate transporter *Slc1a2*, which has been shown to be mutated in epileptic encephalopathies (Epi, 2016).

In conclusion, our study has identified extracellular signals and transcription factors that drive distinct parts of the astrocyte maturation programme through chromatin and transcriptional reorganisation, and thereby regulate important aspects of astrocyte functions in brain homeostasis. With this, our study opens up new directions of research which may contribute to better understand and ultimately treat neurological disorders involving astrocyte dysfunction.

## Supporting information

Supplemental Information (Figures with legends, table legends)

## Acknowledgments

We thank Maria del Mar Masdeu and Lan Chen for technical support, the Science Technology Platforms of the Francis Crick Institute and in particular Amelia Edwards and Deb Jackson from the Advanced Sequencing Facility, Harshil Patel and Nourdine Bah from Bioinformatics & Biostatistics and the Biological Research Facility for their support. This work was supported by a grant from the Medical Research Council to F.G. (10408), by funding to the Francis Crick Institute, by Cancer Research UK (FC0010089), the UK Medical Research Council (FC0010089) and the Wellcome Trust (FC0010089), a research fellowship by the German Research Foundation (DFG) to M.L. (LA 4031/1-1).

## Author Contributions

Conceptualization: M.L., F.G.; Methodology: M.L., R.G., F.G.; Formal Analysis: M.L., R.G.; Investigation: M.L., R.G.; Data Curation: M.L., R.G.; Writing-Original Draft: M.L., F.G.; Writing-Review & Editing: M.L., R.G., F.G.; Visualization: M.L., F.G.; Supervision: F.G.; Project Administration F.G.; Funding Acquisition: M.L., F.G.;

## Declaration of Interests

The authors declare no competing interests.

## STAR Methods

### RESOURCE AVAILABILITY

#### Lead Contact

Requests for further information, resources and reagents should be directed to the Lead Contact, Francois Guillemot.

#### Materials Availability

Plasmids generated are available on request from the Lead contact.

#### Data and Code Availability

Sequencing data generated in this study will be deposited to the NCBI GEO data repository.

### EXPERIMENTAL MODEL AND SUBJECT DETAILS

#### Mouse models

Animal care and procedures were performed in accordance with the guidelines of the Francis Crick Institute, as well as national guidelines and laws, under the UK Home Office Licence PPL PB04755CC. Mice were housed in standard IVC cages with ad libitum access to food and water, under a 12/12h light/dark cycle. Striatal astrocytes were purified from brain tissue of female mice from a mixed background at postnatal day 3 (P3), P7 or at an age of 12-14 weeks (3m). Cortical astrocytes were purified from brain tissue of male C57BL/6Jax mice was used, at P4, or at an age of 6-10 weeks (2m).

#### Primary neural stem/progenitor cells

Primary neural stem/progenitor cells (NSCs) were derived from mice of a mixed MF1-derived background. Three batches were used in this study: batch X6 (from adult subventricular zone tissue, age 8 weeks), batches X8 and X9 (from P3 postnatal cortical grey matter tissue).

Tissue was dissociated with the “Neural Tissue Dissociation Kit (P)” (Miltenyi, 130-092-628) and cell preparations were initially expanded for 1-2 passages as neurospheres in *basal medium* ((DMEM/F-12 + Glutamax (Invitrogen 31331-093) + 1x N2 supplement (R&D Systems, AR009) + 1x Penicillin-Streptomycin (ThermoFischer Scientific, 15140)) supplemented with 20 ng/ml EGF (Preprotech, 315-09-100) + 20 ng/ml FGF2 (Peprotech, 450-33) + 5μg/mL Heparin (Sigma, H3393-50KU).

Subsequently cells were further expanded as adherent cultures in *basal medium* supplemented with EGF + FGF2 + Heparin (as above), + 2 ug/ml Laminin (Sigma, L2020) (*EGF/FGF2-medium*), on plates pre-coated for >1h with EGF/FGF2 medium. For passaging, cells were detached with Accutase (Sigma, A6964). Cells were maintained at 37°C, 5% CO2 and used for experiments at total passage 7-25.

### METHOD DETAILS

#### MACS-based astrocyte purification

Astrocytes were purified using the “Anti-ACSA-2 MicroBead Kit, mouse” (Miltenyi, 130-097-678) with the protocol recommended by the manufacturer for adult astrocytes.

Brains of two animals per preparation were cut into coronal slices and cortical grey matter or striatal tissue (excluding the subventricular zone) was dissected. The tissue was cut to small pieces and dissociated with the “Neural Tissue Dissociation Kit (P)” (Miltenyi, 130-092-628), incubating for a total of 30 min at 37C under gentle agitation. Gentle mechanical dissociation was applied after addition of DNase at 15 min, and more thorough dissociation after 25 min (using fire-polished pipettes). After removal of remaining incompletely dissociated tissue pieces using a 70 um cell strainer, the enzymatic dissociation reaction was stopped with calcium-containing HBSS buffer. All subsequent steps were performed on ice or at 4C with pre-cooled reagents.

The cells were collected by centrifugation (300g, 10 min, 4C) and resuspended in 6.2 ml DPBS + 1.8 ml “Debris removal solution” (Miltenyi 130-109-398). The suspension was overlaid with 4 ml DPBS, followed by centrifugation at 3000g, 10 min, 4C. The upper two gradient layers were removed (retaining ~ 1 ml), 10 ml DPBS added, and the cells collected by centrifugation at 1000g, 10 min, 4C. Cells were incubated in “Red blood cell removal solution” (Miltenyi 130-094-183) for 10 min. After addition of 10 ml PB buffer (DPBS + 0.5% BSA), cells were collected by centrifugation (300g, 10 min, 4C), resuspended in 80 ul PB + 10 ul Fc-Block from the ACSA2-Kit, and incubated for 10 min. 10 ul magnetic-bead conjugated Anti-ACSA2-antibody was added for 15 min. Cells were collected by centrifugation and resuspended in 500 ul PB. The suspension was passed through a 70 um cell strainer and a pre-equilibrated MS column (Miltenyi 130-042-201) in an magnetic separator (OctoMACS, Miltenyi 130-042-109). The column was washed twice with PB before the column was removed from the separator to wash out the purified astrocyte preparation with PB.

#### Astrocyte differentiation in vitro (2D cultures)

Neural progenitors maintained *in vitro* at passage 7-25 were plated in *basal medium* ((DMEM/F-12 + Glutamax (Invitrogen 31331-093) + 1x N2 supplement (R&D Systems, AR009) + 1x Penicillin-Streptomycin (Thermo Fisher Scientific, 15140)), supplemented with 20 ng/ml EGF (Preprotech, 315-09-100) + 20 ng/ml FGF2 (Peprotech, 450-33) + 5μg/mL Heparin (Sigma, H3393-50KU) + 2 ug/ml laminin (Sigma, L2020) (*EGF/FGF2-medium*). Progenitors were plated at a density of 40,000 cells/cm^2^ on cell culture dishes coated for >1h in *EGF/FGF2-medium*. Coverslips for immunolabelling were pre-coated before for >1h with 10 ug/ml laminin. After 1 day the medium was changed to *BMP4-medium* (*basal medium* + 2 ug/ml laminin + 20 ng/ml BMP4 (Biolegend, Cat. No. 595302), which was renewed every 2-3 days for 14 days or as indicated in specific figures. For the study of extrinsic maturation signals, after 7 days the *BMP4-medium* was replaced by *basal medium* (+2 ug/ml laminin) or *FGF2-medium* (*basal medium* + 2 ug/ml laminin + 20 ng/ml FGF2), which also was changed every 2-3 days for further 7 days. All cultured cells were maintained at 37°C, 5% CO2. At the end of the differentiation protocol, cells were either fixed with 4 % PFA for 15 min for immunofluorescence, lysed with Trizol for RNA extraction, or detached with accutase (Sigma, A6964).

Sets of experiments with cultures differentiated independently at different timepoints using different neural progenitor preparations were considered as biological replicates.

#### Astrocyte differentiation in vitro in 3D cultures

For 3D cultures, neural progenitors were initially cultured for 7 days as neurospheres in suspension in *EGF/FGF2-medium* without laminin (as described for 2D cultures) at a density of 120,000 cells/ml. Spheres were collected by centrifugation (100g, 3min) and plated at a density equivalent of 100,000 originally plated cells/well in 24-wells. Cells were plated in a basal membrane extract (BME) gel of EGF/FGF2-medium supplemented with 20% “Cultrex PathClear 3-D Culture Matrix RGF BME” (Bio-Techne, 3445-010-01), in wells pre-coated with EGF/FGF2 + 20% BME. Cells were differentiated with BMP4-medium, followed by *basal medium* or *FGF2-medium* (each supplemented with 2% BME) as described for 2D cultures.

As 3D conditions may result in more heterogenous differentiation, e.g. due to residual growth factors in the 3D gel, or limited diffusion of growth factors into the gel, MACS purification was performed to remove potentially occurring contaminating cells. For this, the gel and the embedded spheres were dissociated by addition of Dispase (Life Technologies, 17105041, final concentration 3 mg/ml) to the culture medium for 1h, and gentle mechanical dissociation. Cells were then collected by centrifugation, resuspended in DPBS + 0.5% BSA, blocked with Fc-Block reagent, labelled with ACSA2 antibodies and purified on MACS columns as described above for the purification of astrocytes from brain tissue.

#### Lentiviral expression of candidate transcription factors in a doxycycline-regulated Tet-on system

*Rorb*, *Dbx2*, *Lhx2*, and *Fezf2* open reading frame (ORF) constructs from Origene (MR225771, MR215327, MR206390, MR207270) were re-cloned into a modified version of the TetO-FUW-EGFP lentiviral expression plasmid (Addgene #30130, (Vierbuchen et al., 2010)). In this vector we had replaced *EGFP* by *Nr1d1* with a C-terminal V5 tag, and added additional restriction sites to facilitate exchange of Nr1d1 with other ORFs. We then replaced *Nr1d1* with *Rorb*, *Dbx2*, *Lhx2* or *Fezf2* to generate V5-tagged doxycycline-inducible lentiviral expression vectors for these transcription factors.

For lentivirus production, these plasmids, the TetO-FUW-EGFP control plasmid, or the FUW-M2rtTA driver construct (Addgene #20342, (Hockemeyer et al., 2008)) were transfected into 293FT cells, together with the third-generation lentiviral packaging/envelop plasmids pMD2.G (Addgene #12259), pRSV-rev (Addgene #12253) and pMDLg/pRRE (Addgene #12251) (Dull et al., 1998). 3 days after transfection, virus particles were collected by ultracentrifugation of the medium. Virus particles were resuspended as highly concentrated virus stock in DPBS, which was stored in aliquots at −80C. Virus titers were determined in BMP4-differentiated astrocytes at day 3, which were infected over-night infection of days with the FUW-M2rtTA and any TetO-FUW virus. Transgene expression was induced by addition of doxycycline (2ug/ml) for 2 days, and determined by immunostaining for EGFP or the V5 tag of the transcription factors.

For the transcription factor overexpression studies, FUW-M2rtTA was transduced along with TetO-FUW-EGFP or any of the transcription factor expression constructs, by over-night infection of astrocytes at a multiplicity of infection (MOI) of 3, at day 6 of the differentiation protocol. After infection, the transgenes were induced by addition of doxycycline to the differentiation medium (BMP4-medium) for 7 days with renewal every 2-3 days.

#### Immunofluorescence stainings and microscopy

Cells on glass coverslips (fixed with 4% PFA and stored in PBS at 4C) were washed with PBS + Triton (0.1%) and incubated with *blocking solution* (10% normal donkey serum in PBS/Triton) for 1 h at room temperature. Subsequently, coverslips were incubated over night at 4C with primary antibodies in *blocking solution*, washed three times with PBS/Triton, and incubated with secondary antibodies in blocking solution for 2h at room temperature. After one wash with PBS/Triton and 2 washes with PBS, the coverslips were stained with DAPI (1 ug/ml) in 0.5x PBS for 15 min, and washed again with 0.5x PBS before mounting. Images were acquired with a Leica SP5 confocal microscope with a 20x objective, with z-steps of 1 um through the whole cell monolayer (approx. 8-15 z-layers).

#### Single cell RNA-sequencing and analysis

For single cell mRNA-Seq, single cell suspensions of striatal astrocytes were prepared by MACS as above. Samples were prepared using the 10x Chromium Next GEM Single Cell 3ʹ v3.1 kit. For each sample, cells were quantified using an automated cell counter and viability assessed via trypan blue staining. Where possible, approximately 10,000 cells were loaded into the 10x Genomics Chromium Controller (sample PN2 (P3) had a low cell count, and 4,000 cells were loaded). Following GEM recovery, cDNA was amplified and final libraries prepared according to the manufactuer’s instructions. Finished libraries were quantified using the Qubit (Thermofisher) and TapeStation (Agilent) and pooled for sequencing with the objective of achieving 50,000 reads per cell. Sequencing was carried out on the HiSeq 4000 (Illumina), with 28 cycles for sequencing read 1, 8 cycles for index read 1, and 98 cycles for sequencing read 2.

The resulting data was demultiplexed using the 10x cellranger mkfastq function according to manufactuer’s instructions. Reads from individual samples were mapped to the mouse genome (mm10) and quantified with the cell ranger count function (v3.0.2, option --expect-cells=3500). The generated count matrices were imported into Seurat (v3.1.1) (Stuart et al., 2019) and aggregated using the Seurat “merge” function. Low quality cells with more than 10% mitochondrial transcripts or less than 500 detected genes were removed, as were cells with more than 4500 genes detected (potential duplicates). Expression values were normalised with the “SCTransform” function, followed by dimension reduction with “RunPCA”, and generation of a UMAP plot using “RunUMAP”. “FindNeighbours”, followed by “FindClusters(…, resolution = 1.5) was used to identify cell clusters. “Dimplot(…, group.by = "orig.ident")” was used to analyse contribution of different samples to the clusters. “FeaturePlot” and “DotPlot” functions were used to plot expression of marker genes to identify the cell type identity of the clusters.

To characterise gene expression changes along the hypothetical astroglial maturation trajectory, transcriptomes of the relevant clusters 0, 5, 6, 7, 17 were isolated, and differentially expressed genes between the clusters identified with the “FindAllMarkers” function. To further analyse the expression changes between the different clusters, the matrix of SCT transformed expression values of these genes was extracted, the mean expression per cluster was calculated and centered on the mean expression of all analysed astroglial clusters.

These normalised expression values were plotted and grouped by hierarchical clustering into 60 initial clusters using the “pheatmap” R package (version 1.0.12), with the parameters “pheatmap(…, clustering_distance_rows = “euclidean”, clustering_method = “ward.D2”, cutree_rows = 60)”. Gene modules with similar profiles were then manually merged to the final 7 clusters shown in Figure 1F.

Gene Ontology analysis was performed using the R package “clusterProfiler” (version 3.12.0) (Yu et al., 2012), using the function “enrichGO(…, OrgDb = org.Mm.eg.db, keyType = ‘SYMBOL’, ont = “BP”, pAdjustMethod = “BH”, pvalueCutoff = 0.01, qvalueCutoff = 0.05)”.

#### Bulk RNA sequencing

RNA was extracted using Trizol, from frozen pellets of MACS-purified astrocytes, or by direct lysis of cultured cells on culture dishes. The “Direct-zol RNA MiniPrep Kit” (Zymo Research, R2052) was used for purification of the extracted RNA, according to manufacturer’s instructions, including the optional Dnase digestion step. The extracted RNA was quantified using the GloMax (Promega) and TapeStation (Agilent) using high sensitivity RNA ScreenTapes, or high sensivity BioAnalyser (Agilent). RNA was prepared into cDNA using the NuGEN Ovation RNA-Seq System V2 kit (Tecan). Following cDNA generation, cDNA was fragmented to an average length of 300 bp using the Covaris E220 focused ultrasonicator (Covaris). Final libraries were then prepared using the NuGEN Ovation Ultralow Library System V2 (Tecan) according to the manufactuer’s instructions. Final libraries were quantified using the TapeStation with D1000 ScreenTapes (Agilent), pooled to 4 nM and sequencing on the HiSeq 4000 system (Illumina) with at least 75 bp single ended reads. The resulting data was demultiplexed using the Illumina bcl2fastq program.

#### RNA sequencing analysis

Reads from RNA-Seq fastq files were aligned to the mm10 genome and quantified using the crickbabs/BABS-RNASeq nextflow pipeline (https://github.com/crickbabs/BABS-RNASeq; git commit id: 335ce47db079d6cc2a7f82f4b762620c4f7f27e2). The pipeline was run using nextflow version 0.30.2 (Di Tommaso et al., 2017) with the command “nextflow run main.nf - params-file params.yml”.

After alignment and transcript quantification, all downstream analysis was performed with R. Raw count normalisation and differential expression analysis was performed using the Bioconductor “DESeq2” package (v1.24.0) (Love et al., 2014). Lowly expressed genes (maximal normalised expression <10) were removed, and genes with an adjusted p-value (padj ≤ 0.05) and at least 2-fold change in expression (abs(log2FC) ≥ 1) were considered to be differentially expressed. Relative expression heatmaps were generated using the “pheatmap” R package (v1.0.12) and plotted as log2 transformed DESeq2 normalised expression values, centered on the mean expression of each gene over all plotted samples. Gene Ontology analysis was performed using the R package “clusterProfiler” (Yu et al., 2012), as described for scRNA-Seq.

Known or predicted transcription factors from the Fantom5 transcription factor database were classified as transcription factors in this study (http://fantom.gsc.riken.jp/5/sstar/Browse_Transcription_Factors_mm9, downloaded 27/10/2018). Published *in vitro* astrocyte gene expression data from other models (Hasel et al., 2017; Tiwari et al., 2018) was re-analysed with the nextflow pipeline above, then merged with the datasets of this study and analysed as above.

#### ATAC sequencing

ATAC sequencing libraries were prepared using the OMNI-ATAC protocol (Corces et al., 2017) with minor modifications:

25,000 MACS-purified astrocytes or dissociated cultured cells were collected by centrifugation (300g, 5 min, 4C), and permeabilized for 3 min on ice in 50 ul ATAC-RSB buffer (Tris/HCl (pH7.4) 10 mM, NaCl 10 mM, MgCl_2_ 3 mM) containing 0.1 % NP-40 (IGEPAL CA-630), 0.1% Tween-20 and 0.01% digitonin. 1 ml ATAC-RSB buffer containing 0.1 % Tween-20 was added and the nuclei were recovered by centrifugation (500g, 10 min, 4C). The pellet was resuspended in 25 ul of transposase reaction mix (12.5 ul 2x TD buffer and 1.25 ul Tn5 transposase (Illumina Cat #FC-121-1030), 8.25 ul PBS, 2.5 ul water, 0.1% Tween-20, 0.01% digitonin), and incubated on a rotational shaker (1000 rpm) for 30 min at 37C to fragment and tag accessible chromatin. DNA was then purified using the “MinElute PCR purification kit” (Qiagen, Cat no. 28004) and eluted in 20 ul EB buffer (Qiagen). 5 ul DNA were were amplified by PCR in 20 ul reactions using NEBNext HiFi 2x PCR Master Mix (New England Biolabs, M0541) and barcoded primers described previously (Buenrostro et al., 2013) for 12 cycles: (1) 5 min 72C, (2) 30s 98C, (3) repeat 12x: 10s 98C, 30s 63C, 60s 72C. The amplified libraries were purified using SPRIselect beads (ratio sample : beads 1 : 1.8) to remove small DNA fragments like primers and primer dimers. For all steps until and including the library amplification step, “Nonstick, RNase-free Microfuge Tubes” (Thermo Fisher, AM12350 or 10676825) were used.

ATAC libraries were quantified using the BioAnalyser (Agilent) and pooled to 4 nM for sequencing on the HiSeq 2500 or HiSeq 4000 system (Illumina). With either instrument, libraries were sequenced with 100 bp paired end reads. Following sequencing, the data was demultiplexed using the bcl2fastq program (Illumina).

#### ATAC sequencing analysis

Reads from ATAC-Seq fastq files were aligned to the mm10 genome, peaks mapped and quantified using the crickbabs/BABS-ATACSeqPE nextflow pipeline (https://github.com/crickbabs/BABS-ATACSeqPE; git commit id: 22edccf72855d42e6692a27385cf50666c8f391c; now superseded; a newer version is available as part of the nf-core project: nf-core/atacseq pipeline (https://doi.org/10.5281/zenodo.2634132 (Ewels et al., 2020)). The pipeline was run using nextflow version 0.30.2 (Di Tommaso et al., 2017) with the command “nextflow run main.nf --design ../design.csv --genome mm10-profile conda --outdir ../results/”. All downstream analysis was performed with R, except transcription factor binding motif enrichment, which was performed using the Homer software (Heinz et al., 2010).

After initial peak calling, annotation and quantification on the merged dataset, the functional peak annotation and classification was extended using multiple published datasets (see also Figure S2B). First, promoters identified by the original pipeline and peaks overlapping with promoters in the regulatory build of the Ensembl database (version 20161111), were classified as “promoters". Other peaks were classified as putative enhancers, if they were overlapping with enhancers it the Ensembl regulatory build, or if they were overlapping with peaks of the enhancer mark H3K4me1 in forebrain P0 or adult cortex in published datasets from the Encode database (https://www.encodeproject.org/; bed files, datasets ENCFF172LKQ, ENCFF746YEV). As a quality control, the ATAC-Seq peaks were also overlapped with DNase-Seq peaks from E18.5 and adult brain (bed files, Encode datasets ENCFF591XUM, ENCFF865BUI). The corresponding bigwig files were downloaded from the Encode database for visualisations in the IGV viewer.

To identify potential regulatory interactions, the original pipeline annotated each peak to the closest TSS. To include potential long-range interactions, distal peaks (i.e. peaks not classified as promoters) were annotated to additional genes, if the peaks were overlapping with regulatory regions for these genes in brain E14.5 or adult cortex described by Shen et al. (Shen et al., 2012), or regulatory regions in adult cortex or cultured neural progenitors (data for confidence level p <0.01) described by Ron et al. (Ron et al., 2017). The conversion of mm9 genome coordinates to mm10 coordinates for these datasets was performed with the Ensembl Assembly Converter (https://www.ensembl.org/Homo_sapiens/Tools/AssemblyConverter?db=core).

Normalisation of read counts and analysis of differential accessibility for the peak regions called on the merged dataset was performed using the Bioconductor “DESeq2” package. Minor peaks with a normalised accessibility < 25 in all samples were removed for further analysis. Peaks with padj ≤ 0.05, abs(log2FC) ≥ 1 were considered as differentially accessible. Bigwig files of the merged replicates of each group (or individual replicates in Figure S2A) were visualised in the IGV viewer with auto-normalisation at a genomic position with a large non-regulated peak (with minimal log2FC determined by DESeq2).

Count matrices for accessibility heatmaps were generated from bigwig files using the R packages “GenomicRanges” (v1.36.1) (Lawrence et al., 2013) and “rtracklayer” (v1.44.4) (Lawrence et al., 2009). The genomic coordinates of the selected peaks were imported from bed files, the bigwig files were imported with the function import(…,format=“BigWig”, as = “RleList”). From these data the counts at peak centers +/− 1 kb were binned in 100 bins of 20bp using “CoverageHeatmap(…, coords = c(−1000, 1000), nbin = 100) and plotted using the “plotHeatmapList” function.

De novo motif analysis in differentially accessible regions was performed on the respective bed files using the Homer findMotifsGenome.pl command (parameters: -size given -mask), with the mm10 mouse genome as background.

### QUANTIFICATION AND STATISTICAL ANALYSIS

#### Quantification of glutamate synthase staining

For quantification of glutamate synthase (GS) stainings (Figure 5B), images of all compared samples were taken in one session with the same settings. Maximum intensity projections of each 2 random areas on 2 coverslips were analysed per independent sample. Intensity of GFP/V5 and GS were measured in regions of interest defined by the DAPI staining using FIJI. For each channel a common intensity threshold for all samples was defined using representative images.

#### Statistical analysis of sequencing data

Statistical analysis of sequencing data was performed with standard methods of the used software tools as outlined in the relevant figures and methods sections.

### KEY RESOURCES TABLE

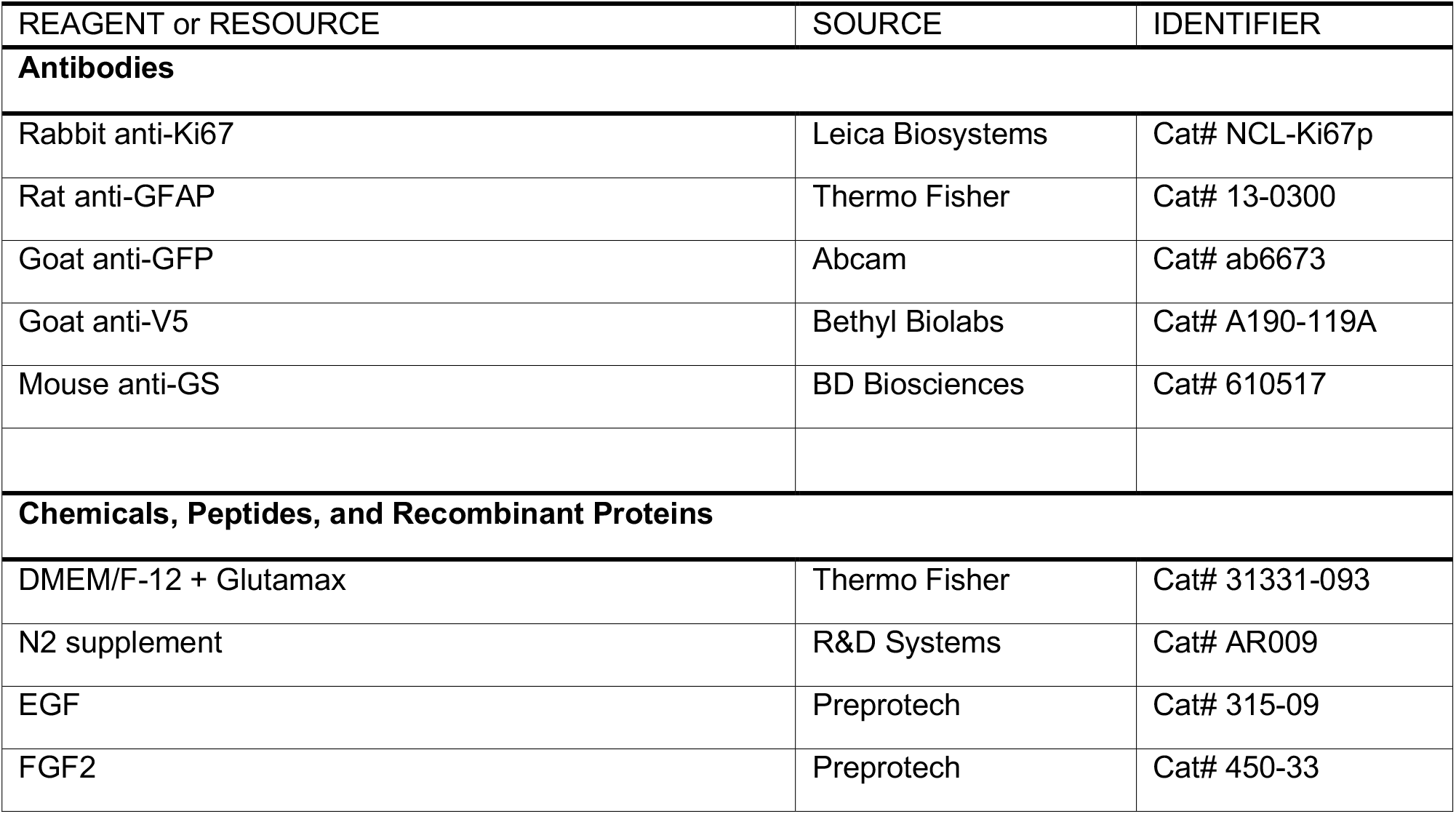
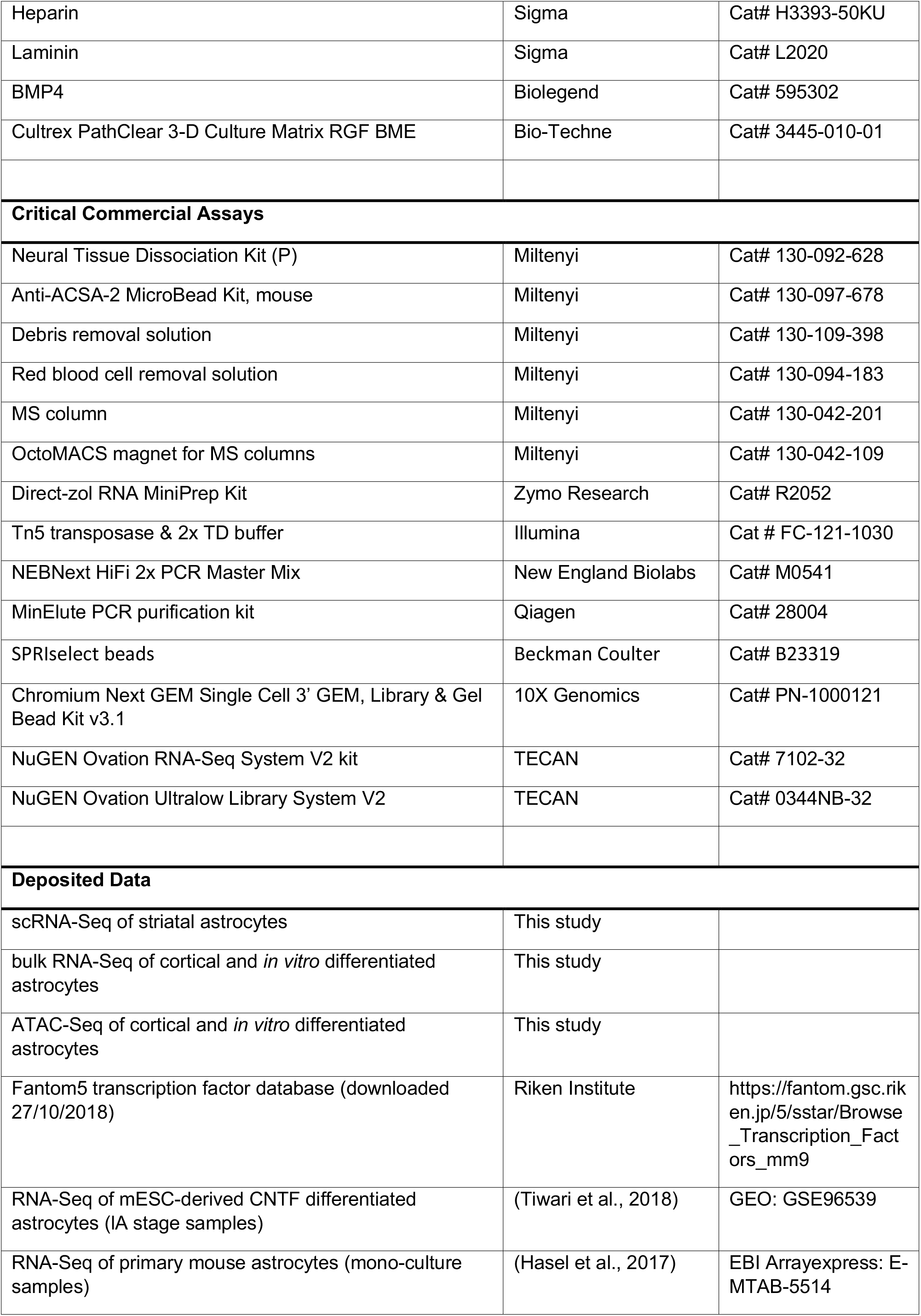
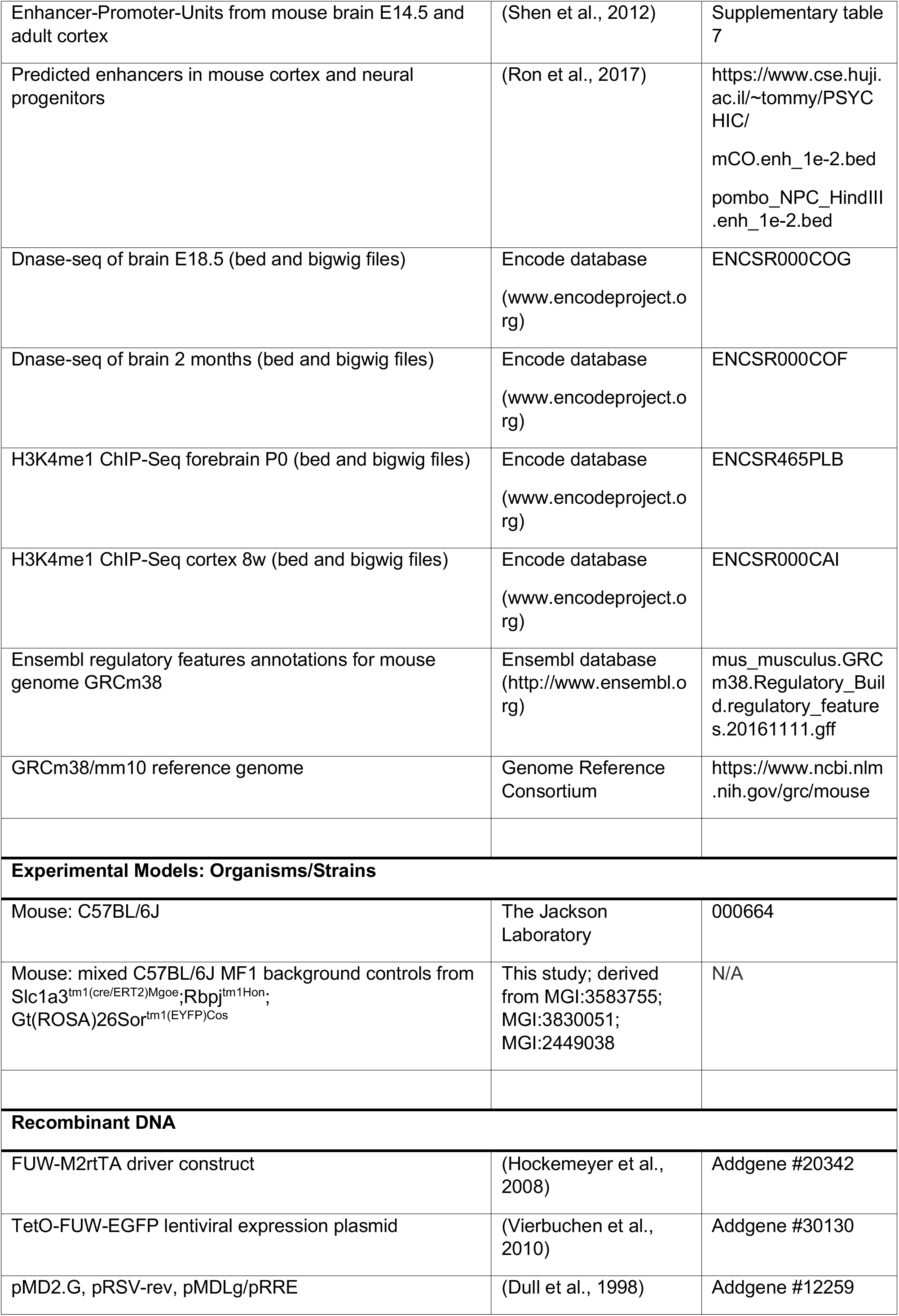
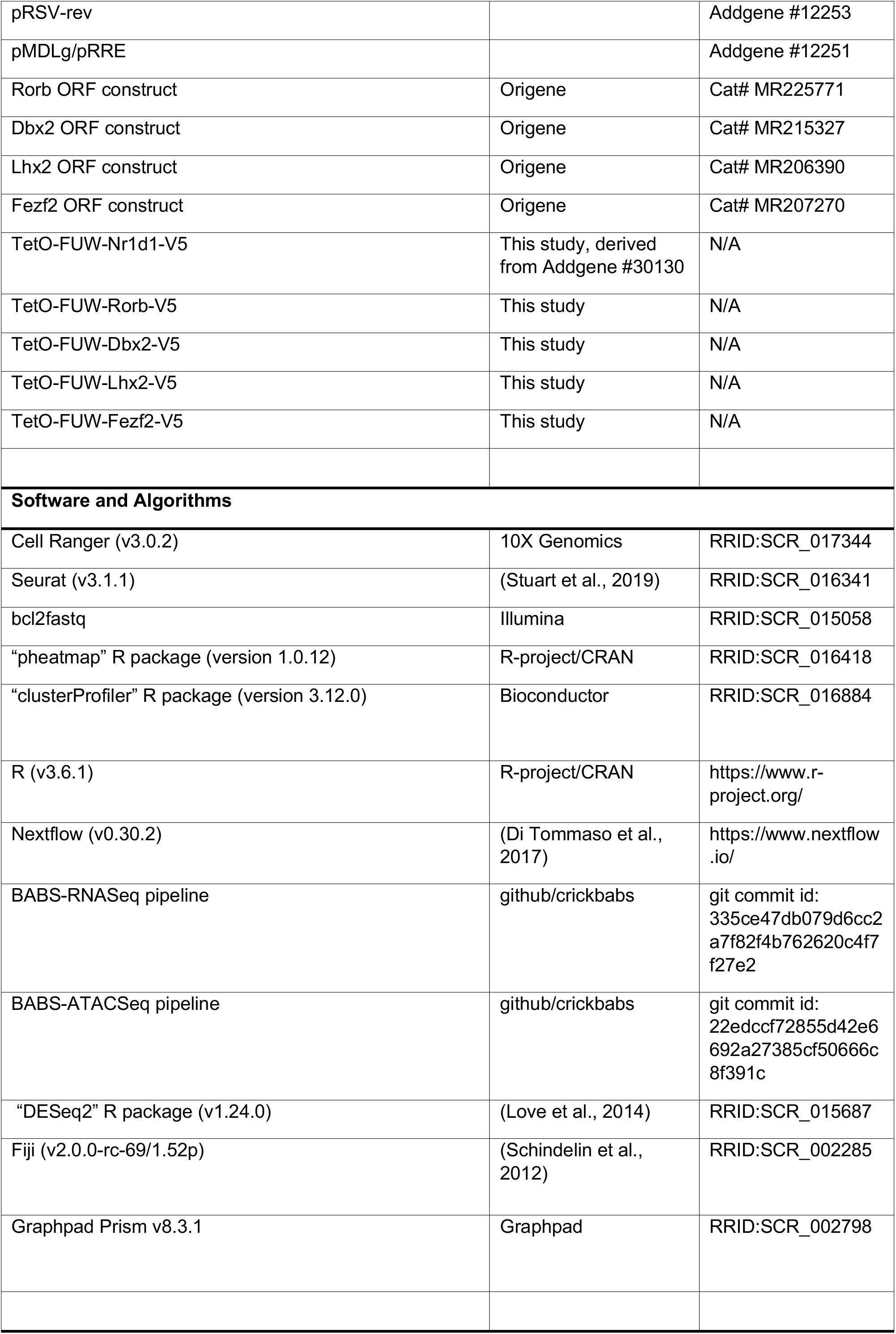

## Notes

### Competing Interest Statement

The authors have declared no competing interest.

### Summary of Updates

Figures and legends included in manuscript text to improve readability

