## Supplemental Information (Figures with legends, table legends) for "Extensive transcriptional and chromatin changes underlie astrocyte maturation *in vivo* and in culture"

**Supplemental Information titles and legends**

**Supplemental Figures**

**Figure S1: Characterisation of cell clusters in striatal sc-RNA-Seq (related to Figure 1)**

**Figure S2: *In vivo* ATAC-Seq peak characterisation (related to Figure 3)**

**Figure S3: Characterisation of astrocytes differentiated *in vitro* from cultured NSCs using BMP4 (related to Fig. 4)**

**Figure S4: Characterisation of transcriptional regulation by Rorb, Dbx2, Lhx2 and Fezf2 expression in cultured astrocytes (related to Fig. 5)**

**Supplemental tables**

**Table S1:** Immature and mature striatal astrocyte genes identified by sc-RNA-Seq; sets of differentially expressed genes with similar expression patterns in striatal astrocytes at P3, P7, 3 months of age; gene ontology terms enriched in immature and mature striatal astrocyte genes (related to Figure 1).

**Table S2**: Genes differentially expressed between cortical astrocytes from P4 and 2-month-old mice identified by bulk RNA-Seq, including common *immature* and *mature* astrocyte genes differentially regulated in both striatal astrocyte sc-RNA-Seq and cortical bulk RNA-Seq (related to Figure 2).

**Table S3:** Annotations of ATAC-Seq peaks detected in combined P4 and 2-month-old cortical astrocytes. Putative types of regulatory elements and linked target genes; overlap with reference datasets (related to Figure 3).

**Table S4:** *Immature* and *mature* genes potentially regulated by differential accessibility of regulatory regions; peak-gene pairs of differentially accessible ATAC peaks and linked *immature* and *mature* genes, with normalized chromatin accessibility and gene expression in cortical astrocytes P4 vs 2 months (related to Figure 3).

**Table S5:** *In vivo* maturation-regulated genes, grouped by their expression in cultured NSCs and BMP4-differentiated astrocytes (related to Figure 4).

**Table S6:** Genes regulated by Rorb, Dbx2, Lhx2 and Fezf2 expression in astrocytes *in vitro*, and associated gene ontology terms; including *mature* genes with low expression in EGFP controls and *immature* genes with high expression; (related to Figure 5).

**Table S7:** Genes regulated by maturation signals (FGF2, 3D culture) in astrocytes *in vitro*, including signal-induced *mature* genes with low expression in controls (related to Figure 6).

**Figure S1**

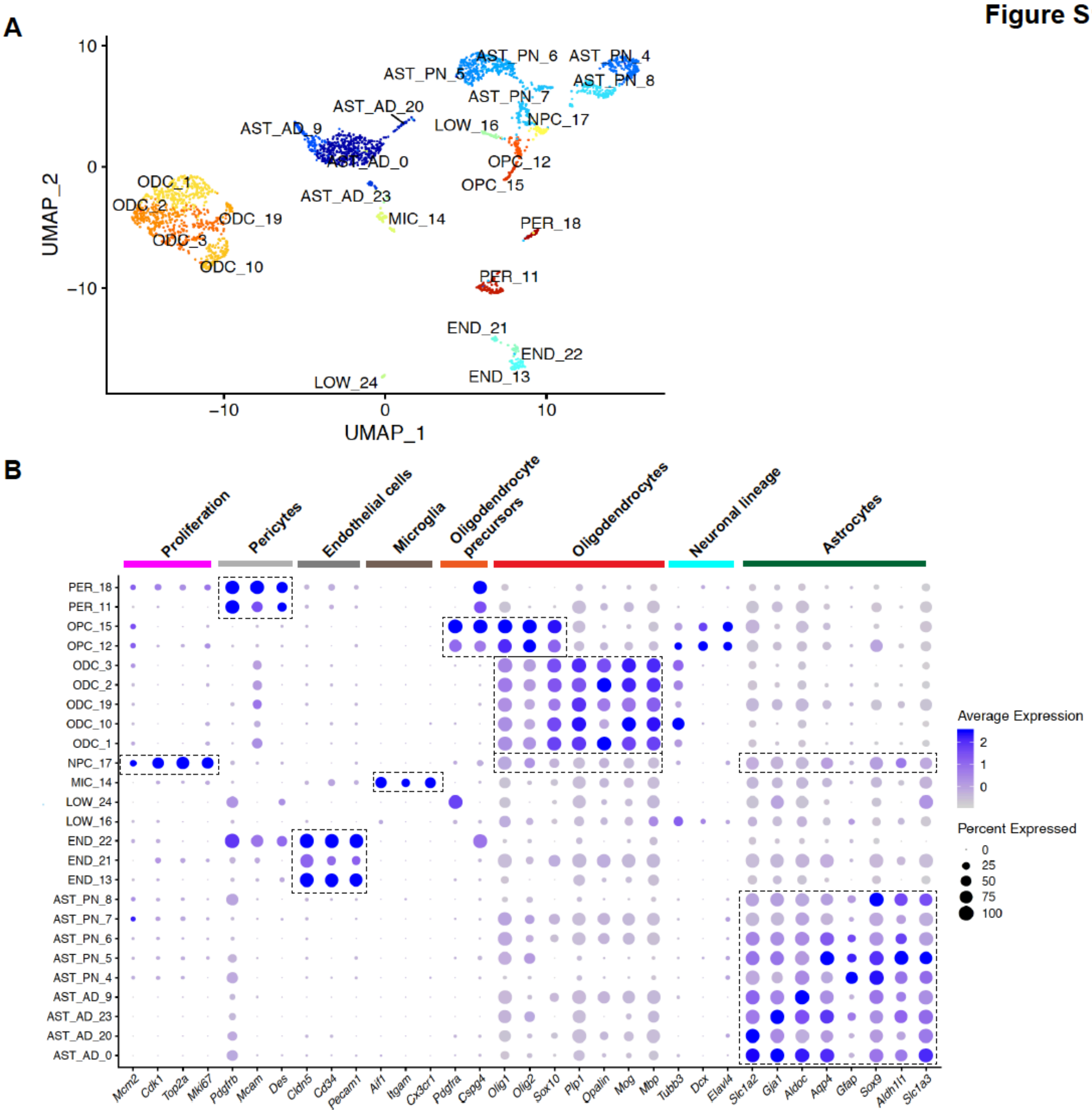


**Figure S1: Characterisation of cell clusters in striatal sc-RNA-Seq (related to Figure 1)**

**(A)** UMAP dimension reduction plots of combined single cell transcriptomes from postnatal and adult striatal astrocyte preparations, showing the cell types corresponding to the clusters, determined by expression of cell type markers shown in (B): AST_PN: postnatal astrocytes (P3/P7), AST_AD: adult astrocytes, NPC: neural (glial) progenitor cells, OPC: oligodendrocyte precursor cells, MIC: microglia, PER: pericytes, END: endothelial cells, ODC: oligodendrocytes, LOW: low marker expression (unidentified).

**(B)** Expression of marker genes used to identify cell types in (A). SCT normalised expression average expression and fraction of cells with detectable transcripts for each cluster.

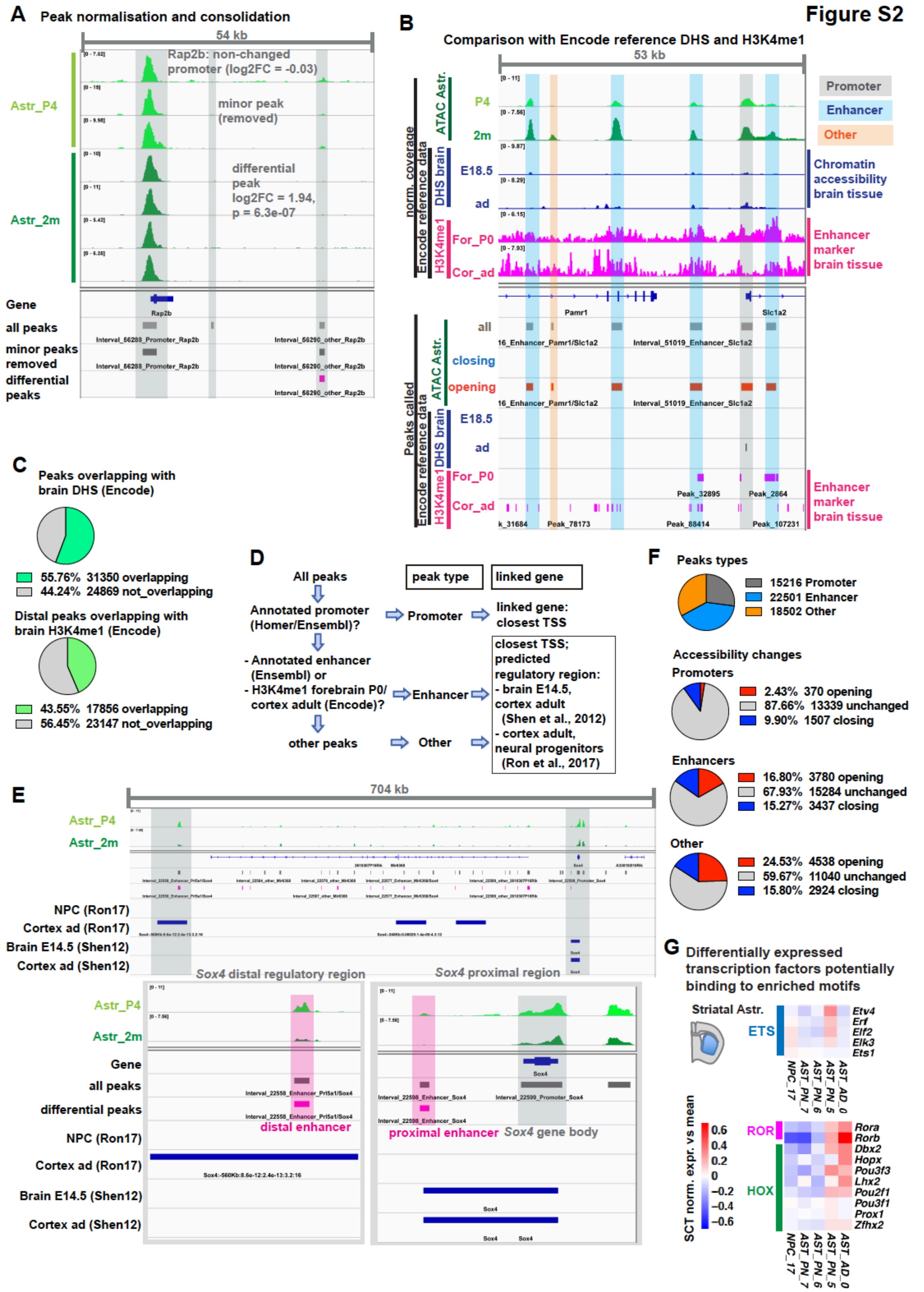


**Figure S2:** ***In vivo*** **ATAC-Seq peak characterisation (related to Figure 3)**

**(A)** Genome tracks show chromatin accessibility (normalised ATAC-Seq read count) in individual astrocyte samples, to demonstrate reproducibility and strategy for peak consolidation and quantification. The *Rap2b* promoter peak is used for normalisation of genome tracks for visualisation. Such large peaks were highly similar between replicates; Example of a minor peak which is removed from the analysis and a differentially accessible peak. The corresponding peak regions called by the analysis pipeline are shown below.

**(B-C)** Comparison of astrocyte ATAC-Seq data with reference datasets from the ENCODE database. DNase-Seq as alternative method to assess chromatin accessibility (whole brain samples E18.5 and adult); ChIP-Seq for the H3K4me1 enhancer mark (tissue samples forebrain P0 and adult cortex); (B) Genome tracks of merged ATAC-Seq data and reference datasets, and corresponding peak regions called by the analysis pipeline (astrocyte samples) or published in the Encode database. (C) Quantification of the overlap of astrocyte ATAC peaks with the reference peak sets. The low signal-to-noise-ratio suggests that the H3K4me1 reference datasets are of relatively poor quality and that the figure of 44% overlap with ATAC peaks is an underestimate.
**(D-E)** Strategy for peak classification based on database annotations and overlap with the enhancer mark H3K4me1 from the reference datasets detailed above. Strategy for the identification of potential target genes of putative enhancers (distal ATAC peaks) located within published regulatory regions. (E) Genome Track of ATAC-Seq data and published regulatory regions around the locus of the *immature* gene *Sox4*, as illustration of this strategy: the highlighted peak in a distal *Sox4* regulatory region may represent a maturation-regulated *Sox4* enhancer that would not be linked to *Sox4* using only a conventional closest TSS approach.
**(F)** Classification of peaks identified in the merged astrocyte dataset based on the approach outlined in (D), and changes in their accessibility during cortical astrocyte maturation from P4 to 2 months of age.
**(G)** Expression of transcription factors in striatal astrocytes that might bind to motifs in differentially accessible regions. scRNA-Seq data (from Figure 1), heatmap of mean relative expression in the astroglial lineage clusters 17,7,6,5,0.
Differential peaks: padj $\leq 0.05$, abs(log2FC) $\geq 1$ (DESeq2 analysis)

Figure S3

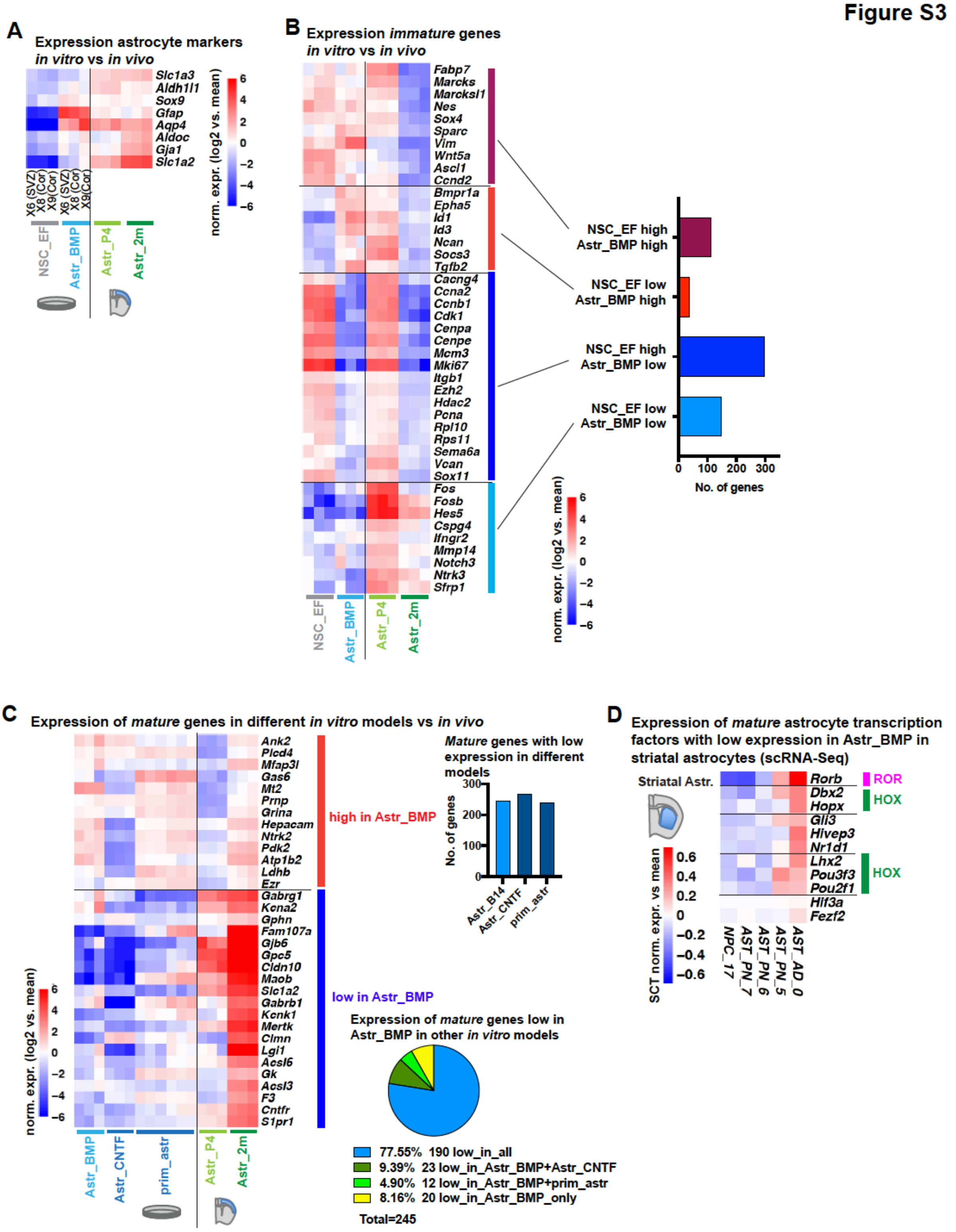


**Figure S3: Characterisation of astrocytes differentiated *in vitro* from cultured NSCs using BMP4 (related to Fig. 4)**

**(A)** Expression of astrocyte markers in astrocytes differentiated *in vitro* with BMP4 for 14 days (Astr_BMP), compared to the original NSCs cultured in EGF/FGF2 medium (NSC_EF), which had been derived from the adult subventricular zone (preparation X6 SVZ) or P3 cortex (preparations X8/9 Cor). RNA-Seq, including cortical astrocytes *in vivo* (Astr_P4 and Astr_2m, from Figure 2).

**(B)** Expression of *immature astrocyte-specific* genes (from Figure 2C) *in vitro* and *in vivo*. Heatmap of selected genes; and total number of genes, grouped by expression grouped by their expression patterns in cultured astrocytes; “NSC_EF/Astr_BMP high” refers to significantly higher expression compared to Astr_2m, “low” to equal or lower expression.

**(C)** Expression of *mature* genes (from Figure 2C) in different *in vitro* astrocyte models. BMP4 differentiated astrocytes from this study (Astr_BMP), astrocytes differentiated from embryonic stem-cell-derived NSCs differentiated with CNTF (Astr_CNTF) and cultured primary astrocytes (prim_astr) from published datasets (Tiwari et al., 2018; Hasel et al., 2017). Heatmap of selected genes, and number of genes with low expression in the different *in vitro* models compared to adult cortical astrocytes.
**(D)** Expression of *mature* transcription factors in striatal astrocytes, which show a low expression *in vitro* (compared to adult cortical astrocytes). scRNA-Seq data (from Figure 1), heatmap of mean relative expression in each of the astroglial lineage clusters 17,7,6,5,0.
Differential genes: padj $\leq 0.05$, abs(log2FC) $\geq 1$ (DESeq2 analysis)

Figure S4

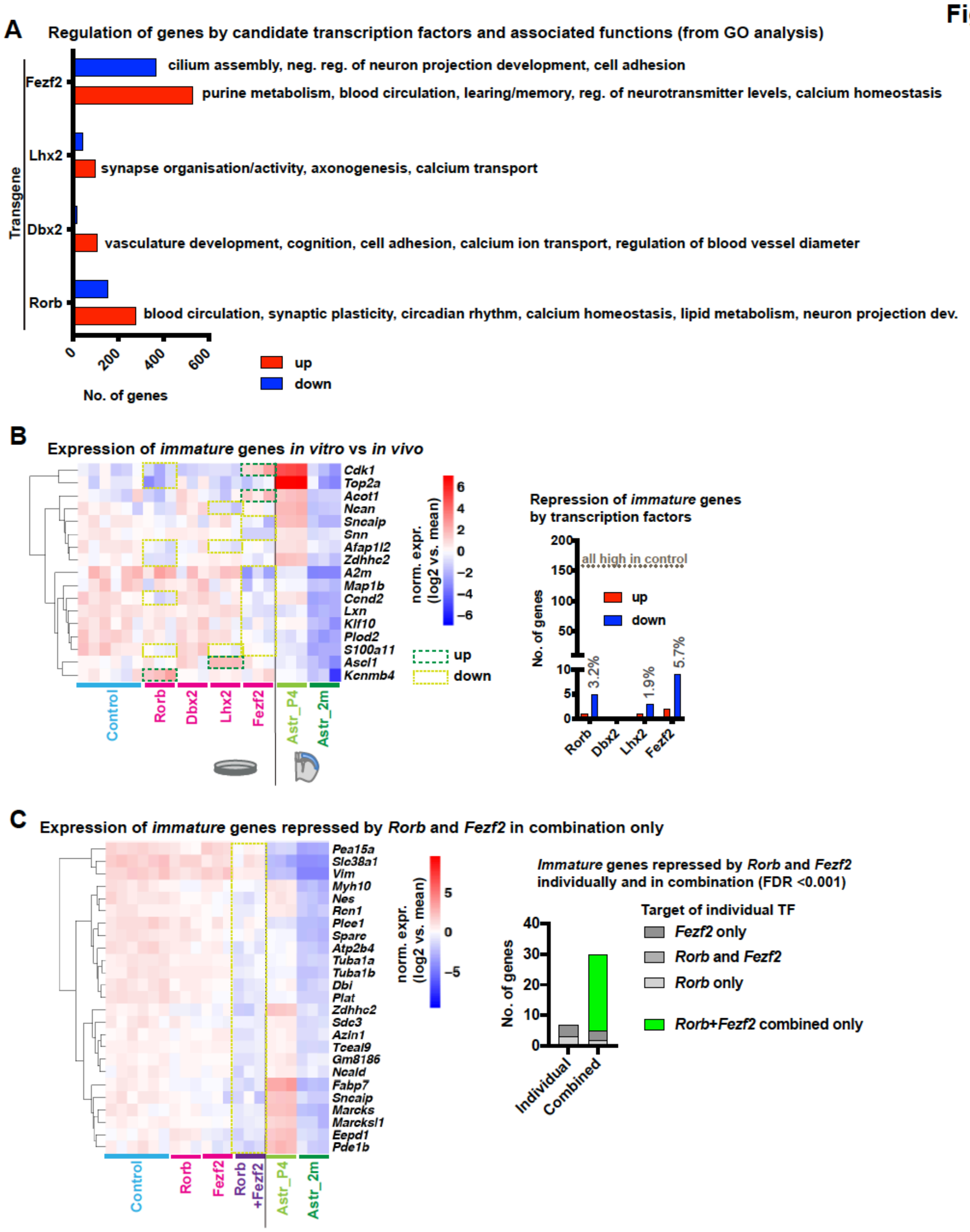


**Figure S4: Characterisation of transcriptional regulation by Rorb, Dbx2, Lhx2 and Fezf2 expression in cultured astrocytes (related to Fig. 5)**

**(A)** Total number of genes differentially expressed in astrocytes expressing candidate transcription factors compared to EGFP controls. Putative functions of these genes from GO analysis (details see Table S7).

**(B)** Expression of *immature astrocyte-specific* genes (from Figure 2C) compared to in cortical astrocytes *in vivo* (from Figure 2). Heatmap of selected genes; barplot showing that of all immature genes with high expression in EGFP controls small subsets are repressed by the candidate transcription factors.

**(C)** Regulation of *immature* genes by combined expression of *Rorb* and *Fezf2*. Samples from (B) were analysed together with astrocytes co-infected with *Rorb* and *Fezf2* viruses. RNA-Seq, including data from Figure 2.

Differential genes: abs(log2FC) ≥1, padj ≤ 0.05 (B) or padj ≤ 0.001 (C) (DESeq2 analysis);
